# 17β-estradiol status alters NMDAR function and antipsychotic-like activity in female rats

**DOI:** 10.1101/2025.02.10.637465

**Authors:** Kimberly M. Holter, McKenna G. Klausner, Mary Hunter Hite, Carson T. Moriarty, Samuel H. Barth, Bethany E. Pierce, Alexandria N. Iannucci, Douglas J. Sheffler, Nicholas D.P. Cosford, Heather A. Bimonte-Nelson, Kimberly F. Raab-Graham, Robert W. Gould

## Abstract

Low 17β-estradiol (E2) in females of reproductive age, and marked E2 decline with menopause, contributes to heightened symptom severity in schizophrenia (i.e. cognitive dysfunction) and diminished response to antipsychotic medications. However, the underlying mechanisms are unknown. *N*-methyl-D-aspartate receptor (NMDAR) hypofunction contributes to the pathophysiology of schizophrenia, yet impact of E2 depletion on NMDAR function is not well characterized. Quantitative electroencephalography (qEEG), specifically gamma power, is a well-established functional readout of cortical activity that is elevated in patients with schizophrenia and is sensitive to alterations in NMDAR function. Using qEEG and touchscreen cognitive assessments, present studies investigated the effects of E2 on NMDAR function by administering MK-801 (NMDAR antagonist) to ovariectomized rats with or without E2 implants (Ovx+E and Ovx, respectively). Ovx rats were more sensitive to MK-801-induced elevations in gamma power and attentional impairments compared to Ovx+E rats. Further investigation revealed these effects were mediated by reduced synaptic GluN2A expression. Consistent with clinical reports, olanzapine (second-generation antipsychotic) was less effective in mitigating MK-801-induced elevations in gamma power in Ovx rats. Lastly, we examined antipsychotic-like activity of a Group II metabotropic glutamate receptor (mGlu_2/3_) positive allosteric modulator (PAM), SBI-0646535, as a novel therapeutic in E2-deprived conditions. SBI-0646535 reversed MK-801-induced elevations in gamma power equally regardless of E2 status. Collectively, these studies established a relationship between E2 deprivation and NMDAR function that is in part GluN2A-dependent, supporting the notion that E2 deprivation increases susceptibility to NMDAR hypofunction. This highlights the need to examine age/hormone-specific factors when considering antipsychotic response and designing novel pharmacotherapies.

## Introduction

Schizophrenia is a complex, multifactorial disorder that presents with strong individual variability in symptom prevalence, severity, and treatment outcome. Various genetic, environmental, and physiological risk factors, including sex-specific factors, contribute to diverse symptom profiles among individuals. In particular, relationships between 17β-estradiol (E2), the most potent form of endogenous estrogen, and schizophrenia symptomology have been established in clinical populations. Female patients have increased severity of psychotic and affective symptoms during the low estrogen phases of the menstrual cycle [1–4]. Some studies even suggest females diagnosed with schizophrenia have lower overall levels of circulating E2 compared to healthy individuals [5, 6]. Lastly, while males and females have their largest peak in diagnoses during their respective late adolescent periods, only females display a second, smaller peak later in life representing over 75% of diagnoses between 45-50 years of age [7]. This age range corresponds with the menopause transition, which is characterized by hormonal changes including markedly fluctuating levels of E2 followed by a profound decrease in overall circulating levels [8].

Relationships between menopause and schizophrenia symptomology have been well documented in the human population. Many primary symptoms including psychotic (hallucinations and delusions), cognitive (working memory and visuospatial memory) and affective (depression and anxiety) symptoms are more severe in postmenopausal patients [7, 9–12]. Additionally, commonly prescribed antipsychotic medications are less effective in postmenopausal patients suggesting a diminished overall therapeutic response [13–16]. Of relevance, estrogens and selective estrogen receptor modulators (SERMs) have displayed efficacy in improving all symptom clusters when administered as an adjunct treatment approach [17–19]. Taken together, this suggests that decreased E2 during menopause may be a risk factor for late onset schizophrenia and reduced treatment response, yet the underlying mechanism contributing to this risk remains unknown.

*N*-methyl-D-aspartate receptor (NMDAR) hypofunction contributes to pathophysiology of schizophrenia, and this is supported by findings that pharmacological blockade of NMDARs induces psychotomimetic-like effects and cognitive dysfunction in humans and animals [20, 21]. Furthermore, increased resting-state and decreased evoked cortical gamma power, as measured via quantitative electroencephalography (qEEG), have been reported in patients with schizophrenia, and these findings correlate with psychotic episodes and cognitive impairment [22–25]. Importantly, gamma power directly reflects cortical excitatory (E) and inhibitory (I) balance and is influenced by NMDAR manipulation particularly on parvalbumin (PV)-containing interneurons. Systemic administration of NMDAR antagonists as well as selective deletion of NMDARs on PV interneurons leads to elevations in resting-state gamma power [26–29]. Thus, present studies used the noncompetitive NMDAR antagonist MK-801 as a pharmacological probe to evaluate how E2 deprivation may influence NMDAR-related cognitive behaviors and brain function (qEEG).

Physiological factors that impact NMDAR expression and function may affect symptom onset and severity in schizophrenia. Postmortem studies in patients with schizophrenia showed reduced NMDAR subunit mRNA and protein expression in regions including the prefrontal cortex (PFC) [30–33]. Furthermore, in preclinical studies using ovariectomy (Ovx) as a model of surgical menopause (depletes ovarian-derived E2, progesterone, and androstenedione creating a blank-slate to allow for systematic evaluation of specific hormones), E2 administration has been shown to rescue decreases in NMDAR subunit expression and binding in the hippocampus [34–38]. Taken together, this suggests that there is a notable interaction between low E2 levels and NMDAR expression and function. Of relevance, previous research has reported that NMDAR subunits differentially affect gamma power. NMDARs are heterotetramers composed of two obligatory GluN1 subunits and two additional GluN2A-D or GluN3 subunits [39]. GluN2A-preferring, but not GluN2B-selective, antagonists lead to increases in gamma power [40, 41].

As related preclinical studies have largely disregarded females and associated hormonal variables, the goal of the present studies was to determine if E2-deprivation, a putative risk factor for schizophrenia, alters cortical NMDAR function in a subunit-specific manner using behavioral, functional, and molecular techniques in Ovx rats. First, using translational touchscreen cognitive tasks, we found that Ovx rats were more sensitive to MK-801-induced impairments on attention, an effect that was restored in Ovx rats administered E2 (Ovx+E).

Importantly, using qEEG, Ovx rats were also more sensitive to changes in gamma power induced by MK-801 and the GluN2A-preferring antagonist PEAQX. This may be attributed to our findings of reduced synaptic expression of GluN2A in the PFC of Ovx rats. Lastly, as changes in gamma power can serve as a valuable biomarker to evaluate therapeutic efficacy, we compared the ability of olanzapine (OLZ) and SBI-0646535 ([42], compound 74), a metabotropic glutamate receptor subtype 2/3 (mGlu_2/3_) positive allosteric modulator (PAM) to mitigate MK-801-induced functional disruptions. mGlu_2/3_ PAMs and agonists have demonstrated antipsychotic-like activity in preclinical models [43, 44], but have not been evaluated in the context of menopause. In line with the human population, OLZ displayed a sedative profile and was less effective in mitigating MK-801-induced increases in gamma power in Ovx rats. In contrast, SBI-0646535 was equally effective in reducing elevations in gamma power in both groups without a sedative profile supporting this as a promising therapeutic mechanism in E2-deprived patient populations. Ultimately, the present studies provide direct support of the relationship between E2 and schizophrenia, and this report is one of few to systematically test the hypothesis that increased schizophrenia onset and severity post-menopause may be mediated by altered cortical NMDAR function.

## Methods and materials

### Animals

Three-month-old female Sprague–Dawley rats were obtained from Envigo (Indianapolis, IN, USA) and individually housed (EEG studies) or pair-housed (cognition and Western blot studies) in opaque cages (8 in × 10 in × 8 in) in temperature-controlled (range: 70–74 °F) colony rooms. All rats for EEG and Western blot studies had *ad libitum* access to food and water and were maintained on a 12/12 hour light/dark cycle. Rats for cognition studies were food restricted to ≥85% of their free-feeding weights for the duration of the studies. All procedures were conducted in accordance with the Wake Forest University School of Medicine Animal Care and Use Committee and complied with the National Institutes of Health Guide for the Care and Use of Laboratory Animals. Animal number for all experiments were determined based on prior studies [27, 45–48].

### Ovariectomy and hormone administration

All studies consisted of 3 experimental groups: ovariectomized (Ovx), Ovx plus E2 (Ovx+E), and ovary-intact (O-I). Under isoflurane anesthesia, dorsolateral incisions were made and ovaries were ligated at the tip of the uterine horn and removed (Ovx and Ovx+E groups). O-I rats underwent a sham surgery whereby only incisions were made under anesthesia. Next, a small incision was cut in the nape of the neck for placement of a 10-mm long Silastic capsule containing E2 and cholesterol (Ovx +E) for continuous hormone delivery for up to a 5-month period [49]. An empty capsule was implanted in Ovx and O-I groups. Capsules were made as previously described [49, 50] (see supplemental material). Rats were individually housed following surgery and treated with buprenorphine (Sigma-Aldrich; 0.03 mg/kg, SC; twice daily x 2 days) and Alloxate (Pivetal; 1 mg/kg, SC; once daily x 3 days). After 48 hours, rats for cognition and Western blot studies were re-housed with their prior cage partner. Rats recovered for a minimum of 10 days prior to beginning studies. Neuroendocrine status for successful Ovx (diestrus-like) and E2 replacement (estrus-like) was confirmed using vaginal cytology (see supplemental material).

### Synaptoneurosome (SN) preparation

Rats (n=18, 6 per experimental group; Ovx, Ovx+E, O-I) were pair-housed and minimally disrupted for 3 months. Following isoflurane anesthesia and rapid decapitation, tissue samples from the right PFC were collected and homogenized in buffer (50 mM Tris, pH 7.35, protease and phosphatase inhibitors [Halt, ThermoFisher]). As previously described [46, 47], tissue was filtered through 100 μm and 5 μm, filters, respectively, to produce SNs. SNs were centrifuged (14,000g, 20 min, 4°C) and the resulting pellet was solubilized in RIPA buffer (150 mM NaCl; 10 mM Tris, pH 7.4; 0.1% SDS; 1% Triton X-100; 1% deoxychoate 5 mM EDTA; Halt). The insoluble fraction of SNs was removed by centrifugation at 14,000g, 20 min, 4°C. The SN soluble fraction’s protein concentration was determined with the Pierce^TM^ BCA Protein Assay Kit (Thermo Scientific).

### Western Blotting

25 μg of protein/sample was separated on a 10% resolving gel by SDS-polyacrylamide gel electrophoresis (PAGE). GluN1 (anti-GluN1; 1:1000, N308/48, NeuroMab), GluN2A (anti-GluN2A; 1:1000, N327/95, NeuroMab), and GluN2B (anti-GluN2B; 1:1000, 244 103, Synaptic Systems) were visualized with aforementioned primary antibodies and allowed to incubate overnight at 4°C. Membranes were subsequently incubated in fluorescence-conjugated secondary antibodies (AF680; AF800; 1:4000; LiCor, Lincoln, NE) and imaged using the Odyssey CLx infrared imaging system. ImageJ software (National Institutes of Health) was used for densitometry analysis. Proteins of interest (GluN1, GluN2A, and GluN2B) were normalized by actin expression and to the O-I condition.

### Drugs

(+) MK-801 hydrogen maleate (Sigma-Aldrich; 0.03–0.18 mg/kg, subcutaneous (SC)) and PEAQX (Hello Bio; 10 and 30 mg/kg, SC), a GluN2A-preferring antagonist with ∼12-fold greater selectivity for GluN2A than GluN2B [51] (doses were selected based on prior behavioral studies [52]), were dissolved in saline. CP-101,606 (Cayman Chemical; 3-30 mg/kg, SC), a GluN2B-selective antagonist (dose range selected based on prior behavioral studies [53, 54]) was dissolved in 10% Tween80 and saline. OLZ (Sigma-Aldrich; 0.3-1.8 mg/kg, intraperitoneal (IP)) was dissolved in 0.2% acetic acid and saline. SBI-0646535 was synthesized by Sanford Burnham Prebys (see [42] for methods) and dissolved in 2.5% ethanol and 5% Tween80 in sterile water. The pH for all compounds was adjusted to ∼7.4 and administered at 1.0-1.5 ml/kg.

### 5-choice Serial Reaction Time Task (5CSRTT)

Rats (Ovx+E n=8, Ovx n=13, O-I n=10) were trained in operant chambers (Lafayette Instruments, Lafayette, IN) 6-7 days per week to perform the 5CSRTT to measure attention; all cognitive testing occurred in the first half of the rodents’ light cycle. Rats first learned to touch an illuminated white square via nosepoke when presented in one of 5 windows spaced horizontally across the computer touchscreen for a 45 mg chocolate sucrose pellet reward (Bio-Serv). Basic training was conducted as previously described [55] and is detailed further in the supplemental material. Briefly, touching the location where the square was illuminated within the limited hold period resulted in illumination of the receptacle light, a 1-second tone, and reward delivery.

Touching the incorrect location within the limited hold period or omitting (no response within the limited hold period) resulted in the illumination of the house light. The duration the stimulus was illuminated and the limited hold time following stimulus presentation in which the rat was allowed to respond decreased from 60 seconds to 1 and 5 seconds, respectively, across training sessions. Training was complete once rats performed ≥ 80% accuracy and ≤20% omissions for three days, and rats then advanced to the variable task.

### Variable 5CSRTT acquisition and testing

In the variable 5CSRTT task, the duration the square was presented varied across trials to manipulate the attentional demand within each session. The illuminated square was displayed in one of the 5 windows in a pseudorandom order; on any given trial, there were 4 possible stimulus durations, ranging between 0.2 seconds (high attentional demand) and 1.4 seconds (low attentional demand). The stimulus duration range was titrated to each individual rat to ensure their performance was ∼70-80% correct and ≤15% omissions (i.e. some rats’ trials were 0.2, 0.4, 0.6, and 0.8 second stimulus durations whereas others had 0.8, 1.0, 1.2, and 1.4 second stimulus durations). Once rats achieved a stable baseline, MK-801 (saline, 0.03-0.18 mg/kg) was administered using a counterbalanced design with a minimum of 3 days between each test day. Further, rats returned to baseline levels prior to the next test session. 5 Ovx rats were excluded from dosing studies due to inability to achieve a stable baseline. Overall % accuracy (correct trials/total trials-omitted trials) and % omission (omitted trials/ total trials) were assessed following MK-801 administration. Additionally, using custom R scripts, the % accuracy and omission were also calculated at each individual stimulus duration. Rats were excluded from analysis at a given stimulus duration if they omitted all trials; if any were excluded, the n is depicted on the bar graph.

### Paired Associates Learning (PAL) Task

Separate groups of rats (Ovx n=10, Ovx+E n=13, and O-I n=10) underwent touchscreen training and acquisition of the PAL task, a measure of visuospatial working memory, as previously described [48, 56].

Briefly, rats learned to associate three different stimuli (spider, flower, or airplane) with a distinct location on the touchscreen in 1-hour sessions (50 maximum trials). In any given trial, two different stimuli (different PAL task; dPAL) were presented, one in the correct location and one in the incorrect location. A response on the stimulus in the correct location resulted in reward delivery whereas an incorrect response resulted in the illumination of the house light for 5 seconds. Following each incorrect response, correction trials (CTs) were initiated in which the same trial was repeated until the rat made a correct response. CTs did not count towards total trials and served as an indirect measure of perseverative behavior. Acquisition was defined as ≥70% correct for 5 out of 7 consecutive days. 3 rats (1 Ovx and 2 Ovx+E) failed to acquire after >80 training sessions and were excluded from the remainder of the studies.

Following acquisition, the effects of MK-801 (saline, 0.03-0.18 mg/kg) were determined. MK-801 was administered 30 minutes prior to the start of the session in a pseudorandom order. There was a minimum of 72 hours between each dose day and rats were confirmed to return to baseline performance before their next test session. Effects of the mGlu_2/3_ PAM, SBI-0646535 (vehicle, 3-30 mg/kg) and OLZ, a second-generation antipsychotic (saline, 0.3-1.8 mg/kg) alone were also assessed (see supplemental material). Percent correct ((correct trials/number of selection trials) X 100)) and percent of total trials that were CTs ((number of CTs/number of selection trials + number of CTs) X 100)) were calculated. Test sessions in which an animal completed <10 trials were excluded from analysis. Mixed effects two-way ANOVAs were used to examine effects of dose and group. If significant, Šídák’s post hoc tests compared each dose to the vehicle condition and each group across doses.

### Electroencephalography

#### EEG surgery

Prior to Ovx, rats (n= 13 O-I, 20 Ovx, 22 Ovx+E) underwent surgery to implant subcutaneous transmitters (HD-S02; Data Sciences International [DSI], Minneapolis, MN, USA), subcranial wires in contact with the dura for EEG recording, and wires in the nuchal muscle for EMG recording as previously described [27, 48, 57]. Rats were individually housed following surgery and treated with Alloxate (Pivetal; 1 mg/kg, SC x 3 days) and Baytril (Elanco US Inc.; 5 mg/kg, SC x 5 days). Rats recovered for a minimum of 7 days prior to the Ovx surgery.

#### EEG Recordings

EEG, EMG, and locomotor activity data were collected as previously described [34, 59, 61]. All recordings were a total of 7 hours beginning at the onset of the light cycle. First, dose-dependent effects of MK-801, PEAQX, CP-101,606, OLZ, or SBI-0646535 (doses administered in a pseudorandom order) were determined alone when administered 2 hours into the light cycle. Next, we examined effects of OLZ (0.56-1.8 mg/kg) or SBI-0646535 (10 and 30 mg/kg) in combination 0.1 mg/kg MK-801. This dose of MK-801 was chosen as it produced similar elevations in gamma power in all groups (see **Fig. 2**). In these combination studies, OLZ or SBI-0646535 was administered 2 hours after recording onset followed by MK-801 administration 30 minutes later.

#### qEEG Spectral Power Analysis

First, 10-second epochs were manually scored by trained observers as wake, rapid eye movement (REM), non-REM (NREM) sleep or artifact using Neuroscore 3.0 Software (DSI) as previously described [34, 59, 61], by a trained scorer blinded to treatment condition (e.g. dose/group). Following sleep staging, a fast Fourier transformation was used to quantify the relative power spectrum, defined as the contribution of oscillatory activity in 1 Hz intervals (0.5-100 Hz) within each 10-second epoch, using aforementioned parameters [27, 48, 57]. Custom MATLAB scripts separated relative power into each sleep/wake state as individual frequencies (0.5–100 Hz) or frequency bands (delta [0.5–4 Hz], theta [4–8 Hz]; we focused on spectral power during awake periods only given the strong wake promoting effects of NMDAR antagonists [58]. To determine pharmacological effects on the full spectral profile, data in 1Hz bins was averaged across a pre-specified bin size and expressed as a percent change from each individual rat’s 90-minute pre-drug baseline; data is displayed as a group mean. The bin size (i.e. 30-90 min post-dosing for MK-801 and 0-300 min for PEAQX) were determined based off perceived peak effects of each compound on high gamma power to avoid any confounding differences in MK-801 and PEAQX metabolism between groups. OLZ and SBI-0646535 were depicted in the 30-150 min post-dosing period, which corresponds with the drug’s respective half-life period following max absorption [42, 59]. Time-course graphs display changes in individual frequency bands for all compounds as the percent change from baseline in 10 min bins across the 7h recording period. Direct group comparisons for the individual frequency bands were determined by averaging each individual’s percent change from baseline during the pre-specified post-dosing period for NMDAR antagonists or the 60-180 min post-dosing period for combination studies with OLZ or SBI-0646535 and MK-801. Lastly, locomotor activity was simultaneously collected, and activity counts were summed into bins corresponding to time points previously described.

### Statistical Analysis

For Western Blot analyses, group comparisons for synaptic GluN1, GluN2A, and GluN2B expression were analyzed using an ordinary one-way ANOVA followed by Tukey’s multiple comparisons test.

For 5-CSRTT, mixed-effects two-way ANOVAs were used to directly compare overall % accuracy and % omission between groups as well as across the 4 stimulus durations in each group. For PAL, percent correct ((correct trials/number of selection trials) X 100)) and percent of total trials that were CTs ((number of CTs/number of selection trials + number of CTs) X 100)) were calculated. Test sessions in which an animal completed <10 trials were excluded from analysis. Mixed effects two-way ANOVAs were used to examine effects of dose and group. If significant, Šídák’s post hoc tests compared each dose to the vehicle condition and each group across doses.

For EEG studies, a mixed-effects two-way ANOVA followed by Dunnett’s post hoc test was used to analyze dose–effect relationships in each group alone for the full power spectrum and within each frequency band over time. A mixed-effects two-way ANOVA followed by Fisher LSD’s post-hoc test was used for direct group comparisons across all frequencies (0.5-100 Hz) following administration of vehicle, 0.18 mg/kg MK-801, 30 mg/kg PEAQX, and 30 mg/kg CP-101,606. For direct group comparisons of the average % change from baseline in the low and high gamma power bands and summed locomotor activity, a mixed-effects two-way ANOVA was used followed by Šídák’s multiple comparisons test. In all cases, significance was defined as p<0.05.

## Results

### Ovx impacts MK-801-induced impairments in the 5CSRTT but not the PAL task

There is growing support for the use of touchscreen cognition as a standardized approach in preclinical research; many rodent tasks have been reverse-translated from human analogs, maximizing validity of this platform [56, 60, 61]. Present studies implemented the 5CSRTT and the PAL task, which are PFC-and hippocampal-dependent tasks, respectively, to evaluate the effects of E2 deprivation on cognitive domains impacted by both schizophrenia and menopause [60, 62–65]. With the known increases in severity of impairments in global cognitive performance in peri-and postmenopausal patients with schizophrenia [63, 66], we hypothesized that E2 deprivation in Ovx rats would exacerbate NMDAR antagonist-induced impairments in both tasks.

Consistent with our predictions, Ovx rats were more sensitive to MK-801-induced impairments on the 5CSRTT. There were no differences between groups in accuracy at baseline (F_2,23_=1.174, p=0.3271; Ovx: 71.76% ± 7.131, Ovx+E: 74.13% ± 6.42, O-I: 78.22% ± 11.80), in part due to individual performance-dependent titration of stimulus durations. However, MK-801 impaired overall accuracy in Ovx and O-I rats, but not Ovx+E rats (**Fig. 1A**). All groups demonstrated significant increases in omissions compared to the vehicle treatment (**Fig. 1B**). Next, to determine how performance was impacted by varying attentional demand, accuracy across stimulus durations (1-4) was evaluated; lower stimulus durations represent a higher attentional demand whereas higher durations represented low attentional demand. O-I rats only showed impairments at the highest tested dose and stimulus duration (**Fig. 1C**). However, as predicted, Ovx rats demonstrated greater impairments with reduced accuracy across multiple stimulus durations and doses (**Fig. 1D**). Importantly, there were no observed impairments in Ovx+E rats (**Fig. 1E**). For direct group comparisons, see **Fig. S1**. Collectively, these data support the conclusion that E2 depletion increased sensitivity to the disrupting effects of MK-801 on attention.

**Figure 1.**
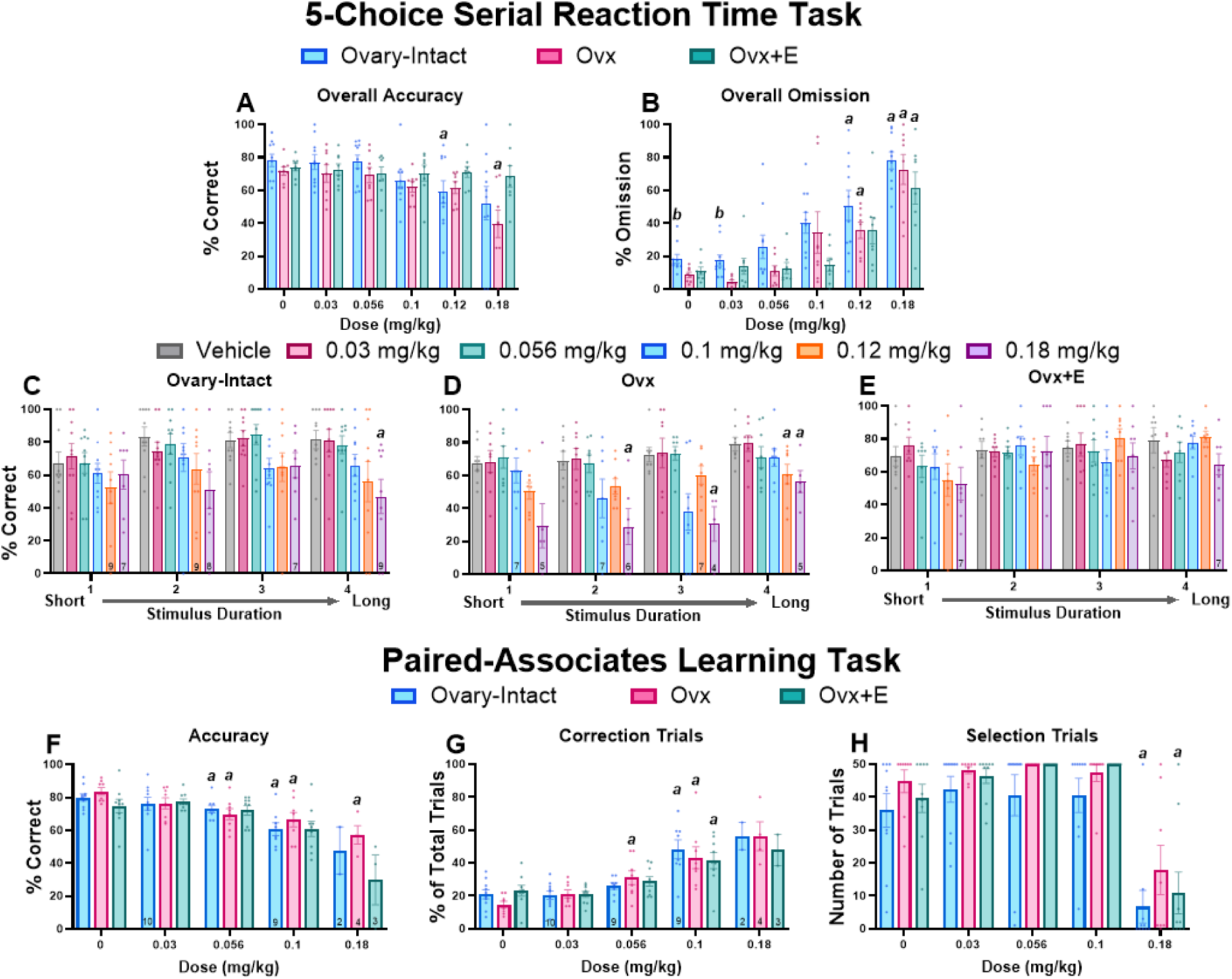
Ovx rats were more sensitive to MK-801-induced impairments on the 5CSRTT but not the PAL task. Bars depict group data as a mean (±SEM). Reductions in overall % accuracy were identified in O-I and Ovx rats in the 5CSRTT (main effect of dose: F_1.908, 43.89_=7.525, p=0.0018) (**A**). Overall % omission was dose-dependently increased in all groups (main effect of dose: F_3.789, 87.15_=48.51, p<0.0001; main effect of group: F_2, 23_=3.994, p=0.0324) (**B**). When separated out by stimulus duration, % accuracy was decreased in O-I (main effect of dose: F_2.630, 132.4_=3.046, p=0.0372; stimulus duration: F_5, 54_=3.667, p=0.0063) (**C**) and Ovx (main effect of dose: F_1.479, 10.36_=5.821, p=0.0265; main effect of stimulus duration: F_2.533, 17.73_=11.14, p=0.0004) (**D**) rats at higher stimulus durations. Ovx+E rats were not affected (ps>0.05) (**E**). In the PAL task, O-I and Ovx rats experienced dose-dependent decreases in overall % accuracy (main effect of dose: F2.167, 39.55=24.11, p<0.0001) (**F**). All groups experienced significant increases in the % of CTs (main effect of dose: F_2.745, 49.42_=49.42, p<0.0001) (**G**). Lastly, the 0.18 mg/kg dose of MK-801 significantly decreased the total number of selection trials in O-I and Ovx+E rats (main effect of dose: F_2.28, 53.11_=44.06, p<0.0001) (**H**). *p*<0.05; *a* compared to the group’s respective vehicle condition; *b* compared to the Ovx group. For 5CSRTT, n=10 O-I, 8 Ovx, and 8 Ovx+E rats; for the PAL task, n= 11 O-I, 8 Ovx, and 9 Ovx+E rats. Some rats were excluded due to omission of all trials within a stimulus duration (**C-E**) or for not performing at inclusion criteria on any given dose day (**G,H**); in those circumstances, the included n is depicted on the bar graph.

In contrast to our predictions, Ovx rats were not more sensitive to MK-801-induced impairments on the PAL task. Interestingly, unlike previous studies using maze-based tasks [67, 68], there were no baseline differences observed in acquisition (prior to MK-801 studies) (data not shown; O-I: 1937 ± 181.7 trials, Ovx: 1797 ± 222.8 trials, Ovx+E: 1809 ± 177.2 trials;F_2,24_= 0.1703, p=0.8444). MK-801 dose-dependently impaired accuracy, but there was no effect of group nor interaction; significant decreases were shown in O-I and Ovx rats (**Fig. 1F**). Additionally, consistent with previous literature in male rats and indicative of perseverative behavior [48, 69], all groups experienced MK-801-induced increases in the % of CTs (**Fig. 1G**). O-I and Ovx+E rats also demonstrated significant reductions in the number of selection trials completed (**Fig. 1H**). Of relevance, several rats from all groups were excluded in accuracy and CT analysis *a priori* as they completed <10 total trials (see reported n’s above bars in **Fig. 1F** and **Fig. 1G**). Overall, results suggest there was no effect of E2 on MK-801-induced working memory impairments at doses that did not produce general decreases in responding. Briefly, neither OLZ nor SBI-0646535 impacted accuracy or CTs, but OLZ significantly decreased selection trials (see supplemental **Fig. S2**).

### MK-801 differentially affected gamma power in Ovx rats

Disruptions in both low and high frequency spectral power have been reported in patients with schizophrenia, as well as following administration of NMDAR antagonists to rodents and humans [23, 26, 70, 71]. Thus, present studies sought to determine if qEEG may reveal NMDAR-related functional differences following E2 deprivation. In conjunction with previous literature showing E2 deprivation can aberrantly influence NMDAR expression, we hypothesized that Ovx rats would demonstrate altered sensitivity to MK-801-induced changes in gamma power.

Consistent with our hypothesis, though there were no group differences at baseline (**Fig. S4**) or following vehicle administration (**Fig. 2A**), decreases in frequencies associated with beta, low gamma, and high gamma were observed in Ovx rats compared to O-I and Ovx+E rats in the 30-90 min post-dosing period of 0.18 mg/kg MK-801 (**Fig. 2B**; for statistics, see **Table 1**). See **Fig. S3** and **Table S1** for dose-related changes across the full spectrum in each group. Further analyses focused on the high gamma power band as the most pronounced group differences were identified in the 50-100 Hz range and gamma has strong relevance for NMDAR manipulations. Direct group comparisons across the MK-801 dose-effect curve within the high gamma power band showed all groups experienced a significantly higher percent change relative to their respective vehicle condition across all tested doses. However, Ovx rats had a blunted percent change from baseline following administration of 0.18 mg/kg compared to Ovx+E animals (**Fig. 2C**). Next, the dose-dependent, timecourse effects of MK-801 on high gamma power were also evaluated within each group (for statistics, see **Table 2**). In O-I rats, all doses elevated gamma power relative to the vehicle condition, and effects of the higher tested doses persisted throughout the 5-hour post-dosing period (**Fig. 2D**). Ovx rats also showed significant increases at all tested doses, yet a unique pattern relative to O-I rats was established.

**Figure 2.**
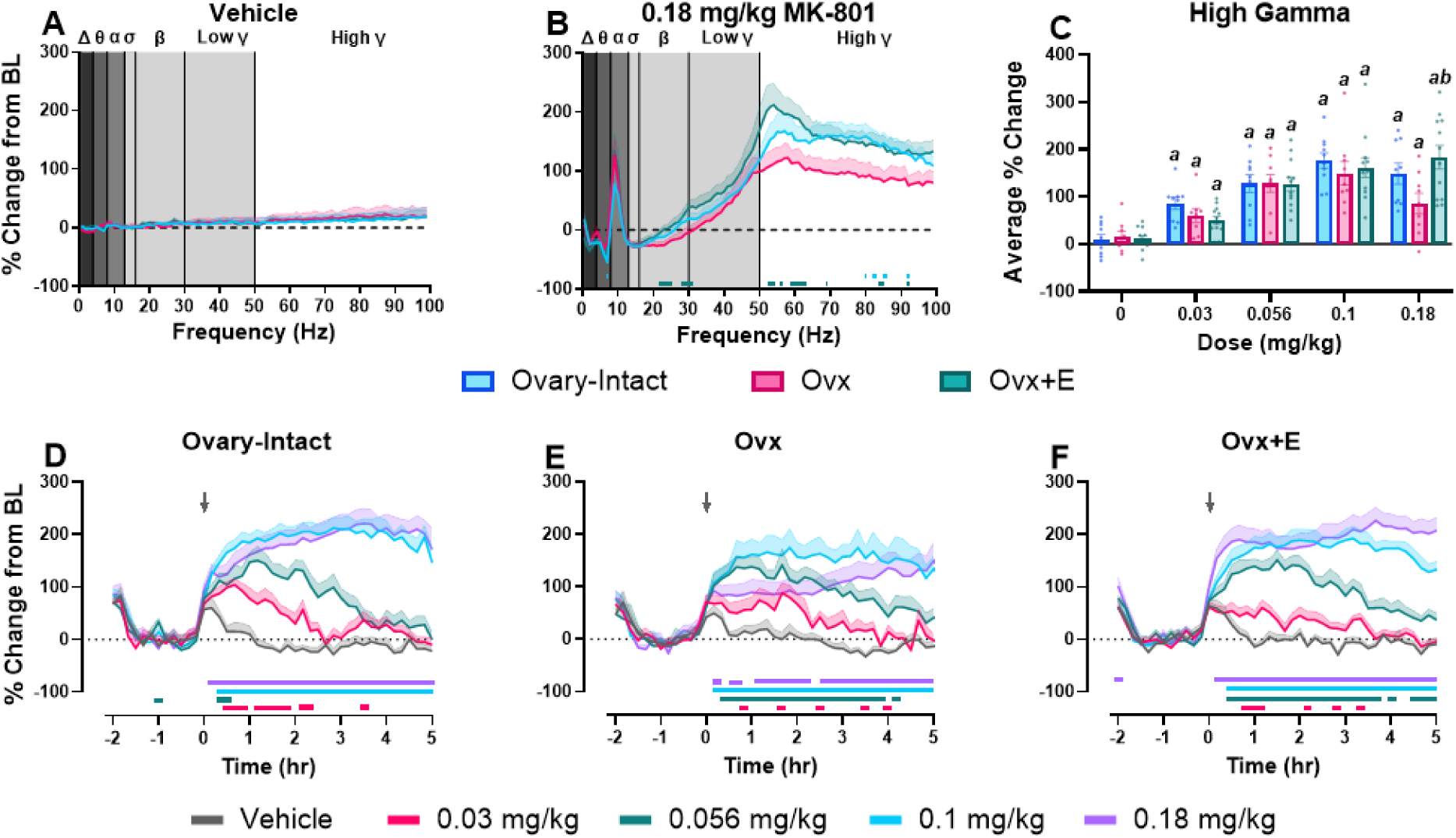
MK-801 differentially affected spectral power in the gamma waveform at the 0.18 mg/kg dose in Ovx rats. All data are shown as a group mean (±SEM). Full spectrum data are presented in 1 Hz bins expressed as the average percent change from baseline during the 30-90 minute post-dosing period. Gray vertical bars represent frequency bands (delta, Δ 0.5–4 Hz; theta, θ 4–8 Hz; alpha, α 8–13 Hz; sigma, σ 13–15 Hz; beta, β 13–30 Hz; low gamma, γ 30–50 Hz; high gamma, γ 50–100 Hz). Group comparisons across the full spectral power range were evaluated following administration of vehicle (**A**) and 0.18 mg/kg of MK-801 (**B**). Then, direct group comparisons evaluated the average % change within the high gamma power band (50-100 Hz) in the 30-90 min post-dosing period at each tested MK-801 dose (main effect of dose: F_2.92, 78.75_ = 46.39, p<0.0001; group X dose interaction: F_8, 108_ = 2.444, p=0.018) (**C**). Lastly, the effects of MK-801 dose on high gamma power over time are displayed as the % change from baseline in 10 min bins across the 7 hour recording period for O-I (**D**), Ovx (**E**), and Ovx+E rats (**F**). p<0.05; Horizontal colored lines corresponding to experimental group represent the 1Hz bins at which O-I or Ovx+E groups were significantly different from the Ovx group (**B**). In **C,** *a* is compared to the group’s respective vehicle condition and *b* is compared to the Ovx group. In **D-F**, horizontal-colored lines matching the respective dose color represent the 10-minute bins at which MK-801-treated groups were significantly different from vehicle-treated groups. n= 9-10 O-I, 8-9 Ovx, and 12 Ovx+E rats.

**Table 1.**
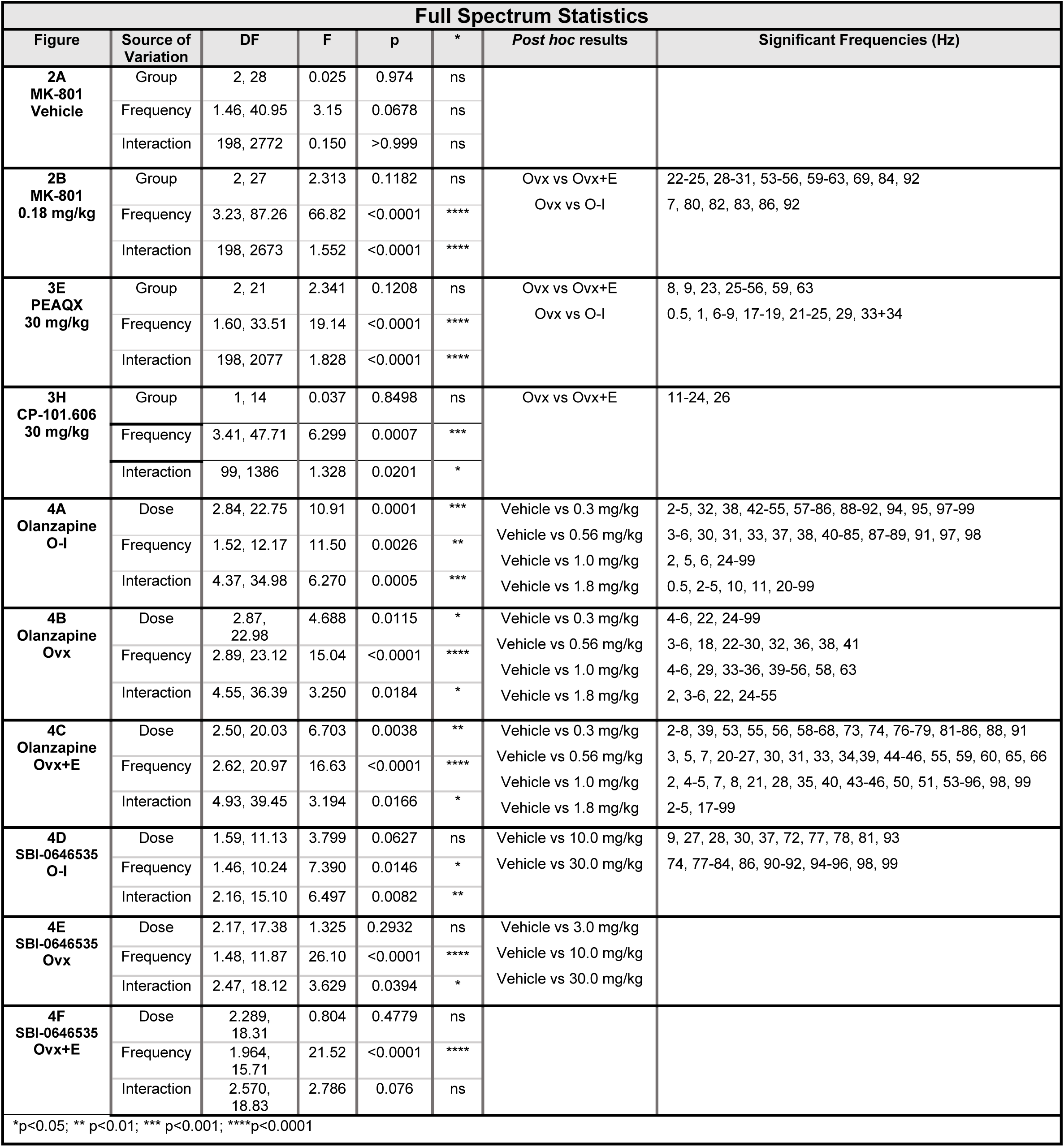
Statistics describing outcomes of MK-801, OLZ, and SBI-0646535 on spectral distribution.

**Table 2.** Statistics describing MK-801 and respective combinations on high gamma power over time.

| Timecourse Statistics |  |  |  |  |  |  |  |
| --- | --- | --- | --- | --- | --- | --- | --- |
| Figure | Source of Variation | DF | F | p | * | Post hoc results | Significant Time points (10-minute bin) |
| MK-801 |  |  |  |  |  |  |  |
| 2D<br>O-I | Dose | 2.50, 22.50 | 33.22 | <0.0001 | **** | Vehicle vs 0.03 mg/kg | 30-50, 70-110, 130, 140, 210 |
|  | Time | 4.15, 37.32 | 46.37 | <0.0001 | **** | Vehicle vs 0.056 mg/kg | -60, 20, 30, 50-200 |
|  | Interaction | 6.31, 51.56 | 13.29 | <0.0001 | **** | Vehicle vs 0.1 mg/kg<br>Vehicle vs 0.18 mg/kg | 20-300<br>10-300 |
| 2E<br>Ovx | Dose | 2.34, 18.71 | 14.29 | 0.0001 | *** | Vehicle vs 0.03 mg/kg | 50, 100, 150, 210, 240 |
|  | Time | 2.71, 21.64 | 16.70 | <0.0001 | **** | Vehicle vs 0.056 mg/kg | 10-250 |
|  | Interaction | 4.82, 36.84 | 6.58 | 0.0002 | *** | Vehicle vs 0.1 mg/kg<br>Vehicle vs 0.18 mg/kg | 10-360<br>10, 30, 40, 70-130, 150-360 |
| 2F<br>Ovx+E | Dose | 2.18, 23.99 | 43.47 | <0.0001 | **** | Vehicle vs 0.03 mg/kg | 50-70, 130, 170, 200 |
|  | Time | 4.19, 46.10 | 48.39 | <0.0001 | **** | Vehicle vs 0.056 mg/kg | 30-220, 240, 260-290 |
|  | Interaction | 7.66, 83.70 | 13.84 | <0.0001 | **** | Vehicle vs 0.1 mg/kg<br>Vehicle vs 0.18 mg/kg | 30-300<br>10-300 |
| Olanzapine + 0.1 mg/kg MK-801 |  |  |  |  |  |  |  |
| 5A<br>O-I | Dose | 2.33, 16.32 | 10.35 | 0.0009 | *** | Vehicle vs 0.56 mg/kg | 0-90, 110, 150, 190 |
|  | Time | 4.14, 28.96 | 67.65 | <0.0001 | **** | Vehicle vs 1.0 mg/kg | 0-160 |
|  | Interaction | 4.90, 33.34 | 5.05 | 0.0016 | ** | Vehicle vs 1.8 mg/kg | 0-160, 200 |
| 5B<br>Ovx | Dose | 2.20, 15.36 | 3.994 | 0.037 | * | Vehicle vs 0.56 mg/kg | 10 |
|  | Time | 2.96, 20.68 | 33.36 | <0.0001 | **** | Vehicle vs 1.0 mg/kg | 10, 30, 50, 60, 80, 150 |
|  | Interaction | 4.28, 29.79 | 2.572 | 0.0548 | ns | Vehicle vs 1.8 mg/kg | 10, 30-80, 100 |
| 5C<br>Ovx+E | Dose | 2.39, 16.75 | 13.43 | 0.0002 | *** | Vehicle vs 0.56 mg/kg | -30, 40, 60-90 |
|  | Time | 3.62, 25.31 | 48.50 | <0.0001 | **** | Vehicle vs 1.0 mg/kg | 30, 50-120, 290 |
|  | Interaction | 5.80, 39.52 | 4.556 | 0.0015 | ** | Vehicle vs 1.8 mg/kg | -30, 10-180, 220, 240 |
| SBI-0646535 + 0.1 mg/kg MK-801 |  |  |  |  |  |  |  |
| 5E<br>O-I | Dose | 1.48, 10.37 | 3.093 | 0.098 | ns | Vehicle vs 30 mg/kg | -80, 60-120, 140 |
|  | Time | 2.85, 19.98 | 54.56 | <0.0001 | **** |  |  |
|  | Interaction | 3.67, 25.08 | 3.777 | 0.0175 | * |  |  |
| 5F<br>Ovx | Dose | 1.15, 8.04 | 9.093 | 0.0147 | * | Vehicle vs 10 mg/kg | -90 |
|  | Time | 1.47, 10.27 | 17.39 | 0.0009 | *** | Vehicle vs 30 mg/kg | 40-210, 230, 250, 290 |
|  | Interaction | 3.50, 22.56 | 5.365 | 0.0045 | ** |  |  |
| 5G<br>Ovx+E | Dose | 1.96, 13.71 | 7.309 | 0.0071 | ** | Vehicle vs 10 mg/kg | -70, -30 |
|  | Time | 1.71, 11.97 | 13.92 | 0.001 | ** | Vehicle vs 30 mg/kg | 50-120, 150, 220, 230, 300 |
|  | Interaction | 4.40, 30.25 | 3.527 | 0.0153 | * |  |  |
| *p<0.05; ** p<0.01; *** p<0.001; ****p<0.0001 |  |  |  |  |  |  |  |

Administration of 0.18 mg/kg to Ovx rats resulted in delayed elevations followed by gradual increases, likely attributed to drug metabolism, that were still observed at the end of the 5 hour recording period (**Fig. 2E**). In contrast, the dose-response profile for Ovx+E rats was comparable to the O-I rats (**Fig. 2F**). Timecourse graphs and direct group comparisons for all frequency bands below 50 Hz are shown in **Figures S5 and S6** and **Table S2**. Briefly, effects of estrous cycle on high gamma power and locomotor activity at vehicle and 0.18 mg/kg MK-801 were evaluated, and no significant effects were identified (**Fig. S7**). Altogether, differences in MK-801-induced changes in cortical gamma power suggest Ovx rats are more sensitive to MK-801 via a leftward-shift in the previously-reported biphasic MK-801 dose-response curve [26, 27]. The comparable increases to other groups identified at low to moderate doses (0.03-0.1 mg/kg) tapered at the highest dose (0.18 mg/kg) in Ovx rats. Ultimately, this pairs well with findings of greater impairments in Ovx rats in the PFC-dependent 5CSRTT as well-established relationships between cognitive performance and gamma power have been reported in preclinical and clinical studies [72].

### Ovx decreases synaptic expression of GluN2A and GluN2B in the PFC and increases sensitivity to PEAQX-induced elevations in gamma power

Next, we evaluated synaptic NMDAR subunit protein expression in the PFC with a specific focus on GluN2A given the known GluN2A-specific effects on cortical gamma power [26, 28]. We thus hypothesized Ovx rats would have lower synaptic GluN2A expression in the PFC. As hypothesized, GluN2A expression was significantly reduced Ovx rats compared to O-I and Ovx+E rats (**Fig. 3A and D**). Lastly, GluN2B expression was also significantly reduced in Ovx rats compared to Ovx+E rats (**Fig. 3B and D**); this finding is consistent with prior literature showing E2 deprivation decreases GluN2B expression in the hippocampus [34, 35, 37, 73]. However, our studies are the first to show E2-mediated changes in synaptic GluN2A expression in the PFC. Importantly, neither Ovx nor Ovx+E rats demonstrated altered synaptic expression of GluN1, the obligatory NMDAR subunit. This suggests decreases in synaptic expression of other NMDAR subunits were likely not attributed to reductions in GluN1 (**Fig. 3C and D**). Full blots are included in **Figure S8**.

**Figure 3.**
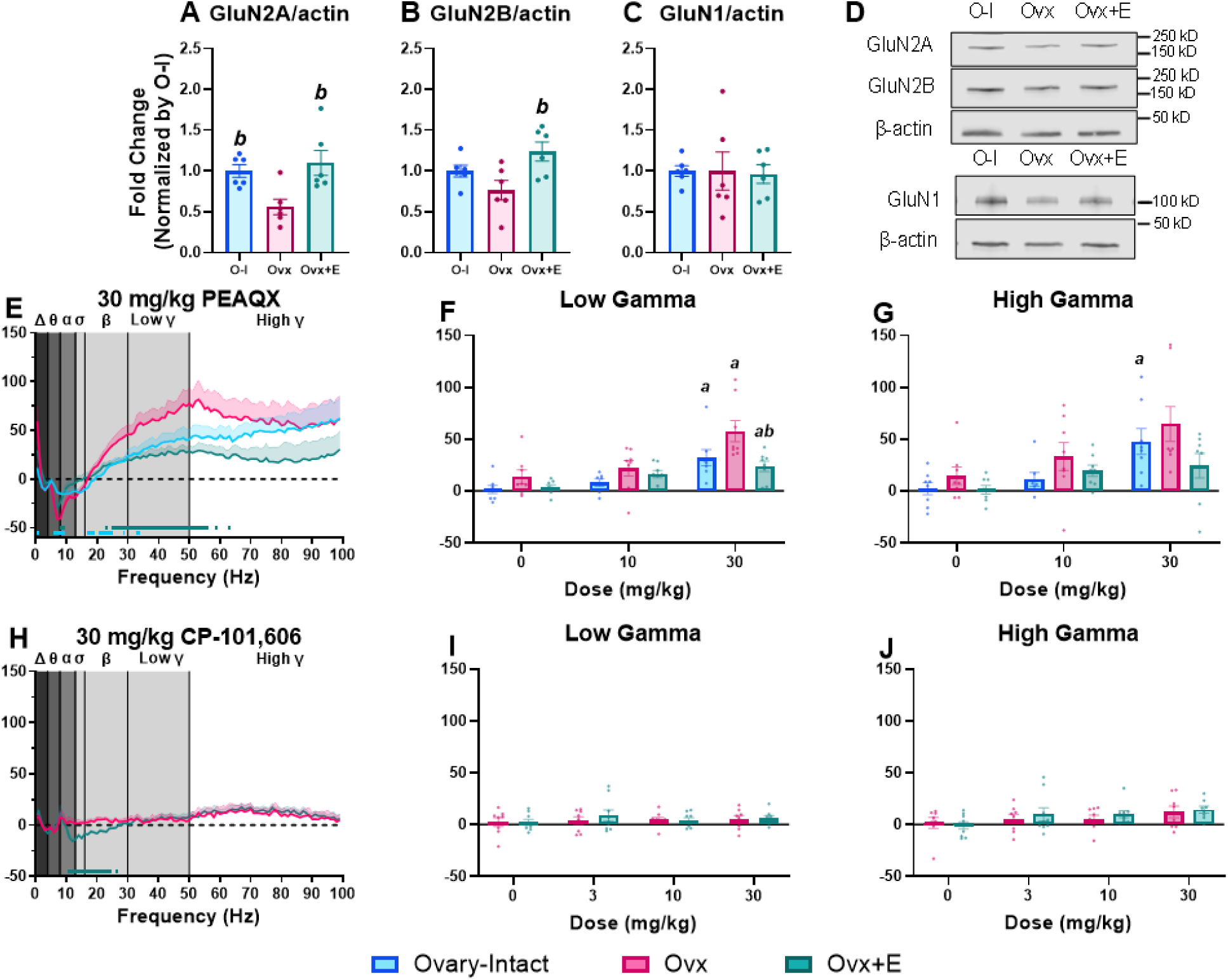
Ovx rats had lower GluN2A and GluN2B synaptic expression in the PFC and were more sensitive to PEAQX-induced elevations in gamma power relative to Ovx+E rats. Densitometry for all data points (1 point per mouse PFC sample) is included; all data are normalized to the O-I condition and expressed as fold change (±SEM); (**A-C**). Western blot analyses of GluN2A expression (F_2,15_= 6.476, p=0.0094) (**A**) and GluN2B expression (F_2,15_= 5.054, p=0.021) (**B**) was significantly lower in Ovx rats (GluN2A: 0.561±0.094; GluN2B: 0.765±0.120) relative to Ovx+E rats (GluN2A: 1.101±0.154; GluN2B: 1.239±0.118). GluN2A expression was also lower in Ovx relative to Ovary-Intact rats (GluN2A: 1.0±0.075; GluN2B: 1.0±0.072). Importantly, GluN1 expression was not significantly different between groups (F_2, 15_ =0.0189, p=0.9813; O-I=1.0±0.066, Ovx= 1.0±0.023, Ovx+E= 1.125±0.114) (**C**). Representative Western blots for GluN2A, GluN2B, and β-actin (above) and GluN1 and β-actin (below) (**D**). In **E-J**, all data are shown as a group mean (±SEM). Full spectrum data are presented in 1 Hz bins expressed as the average percent change from during the 5-hour post-dosing period. Gray vertical bars represent frequency bands (delta, Δ 0.5–4 Hz; theta, θ 4–8 Hz; alpha, α 8–13 Hz; sigma, σ 13–15 Hz; beta, β 13–30 Hz; low gamma, γ 30–50 Hz; high gamma, γ 50–100 Hz). Group comparisons across the full spectral power range were evaluated following administration of 30 mg/kg PEAQX or CP-101,606 (**E,H**). Then, direct group comparisons were assessed for the average % change within the low gamma (main effect of dose: F_1.60,32.87_= 23.4, p<0.0001; group: F_2,21_= 6.686, p=0.0057) (**F**) and high gamma power band (main effect of dose: F_1.568, 32.15_= 13.75, p=0.0001) (**G**) for PEAQX in the 5-hour post-dosing period at each tested dose; CP-101,606 produced no significant effects in the low or high gamma power bands (ps>0.05) (**I,J**). For bar graphs, p<0.05; *a*, compared to the group’s respective vehicle condition; *b*, compared to the Ovx group. Horizontal colored lines matching the respective experimental condition represent the 1Hz bins at which O-I or Ovx+E rats were significantly different from the Ovx condition (**E,H**). In **A-C**, n=6 rats per condition. n= 7- 8 O-I (**E-G**), 8-9 Ovx, and 8-9 Ovx+E rats (**E-J**).

To determine if differences in gamma power between groups following MK-801 administration were GluN2A-mediated, we next examined the effects of PEAQX, a GluN2A-preferring antagonist. Alongside our findings of reduced GluN2A expression, we hypothesized Ovx rats would be more sensitive to PEAQX-induced changes in gamma power. Direct group comparisons at the 30 mg/kg dose showed Ovx rats had a significantly lower percent change in frequencies associated with the theta and alpha bands and a significantly higher percent change in frequencies associated with the delta, low gamma, and high gamma bands compared to Ovx+E and O-I rats (**Fig. 3E**, for statistics see **Table 1**). Interestingly, most differences were in frequencies associated with low gamma. Focusing on the low gamma power band, Ovx rats showed significantly greater elevations following administration of 30mg/kg PEAQX relative to Ovx+E rats (**Fig. 3F**). In the high gamma power band, only O-I rats experienced significant elevations at 30 mg/kg PEAQX (**Fig. 3G**). As expected, CP- 101,606 produced no significant effect on low or high gamma power in Ovx and Ovx+E rats (**Fig 3H-J**). See **Fig. S9 and S10** and **Table S2** for PEAQX and CP-101,606 timecourse for each frequency band. Collectively, these findings suggest that E2 deprivation alters NMDAR expression and function when probed by an NMDAR antagonist and that these disruptions are, in part, a result of altered GluN2A expression.

### SBI-0646535 has a unique spectral profile compared to OLZ

Previous literature shows the efficacy of current antipsychotic medications is blunted in postmenopausal patients with schizophrenia [13, 16]. Thus, it is imperative to investigate novel therapeutic mechanisms that demonstrate greater efficacy in these populations. In line with this previous clinical literature, we wanted to examine the functional response of O-I, Ovx, and Ovx+E rats to OLZ as well as to a compound with a potentially more favorable mechanism, the mGlu_2/3_ PAM SBI-0646535. Ideally, allosteric modulators such as SBI-0646535 can fine-tune this aberrant glutamate efflux associated with NMDAR hypofunction and would ultimately be more effective in E2-deprived conditions assuming this condition has greater glutamatergic disruptions. First, we examined the dose-dependent effects of OLZ and SBI-0646535 alone on the full qEEG spectral profile. Reflective of a sedative profile, all groups showed a significant increase relative to the vehicle condition in low frequencies associated with the delta and theta power bands and a decrease in high frequencies associated with the beta, low gamma, and high gamma frequency bands at all tested doses of OLZ (**Fig 4A-C**; for statistics, see **Table 1**). Interestingly, unlike O-I and Ovx+E rats where nearly all frequencies in the 50-100 Hz range were significantly reduced by 1.8 mg/kg OLZ, Ovx only showed significant reductions below 60 Hz. The broader range of decreases was instead observed at the 0.3 mg/kg dose in Ovx rats, suggesting altered sensitivity to OLZ in the Ovx condition. Alternatively, SBI-0646535 displayed a unique spectral profile compared to OLZ in all three groups. Only O-I rats displayed a significant dose x frequency interaction and, in contrast to OLZ, significant increases in spectral frequencies associated with the high gamma frequency band (>70Hz) were present (**Fig. 4D-F**, for statistics see **Table 1**). The timecourse graphs for all individual frequency bands are displayed in **Fig. S11 and S12** and **Table S3**. Overall, the lack of a sedative qEEG profile of SBI-0646535 alone across groups supports the therapeutic potential of this mechanism without the aberrant side effects reported with current antipsychotic medications.

**Figure 4.**
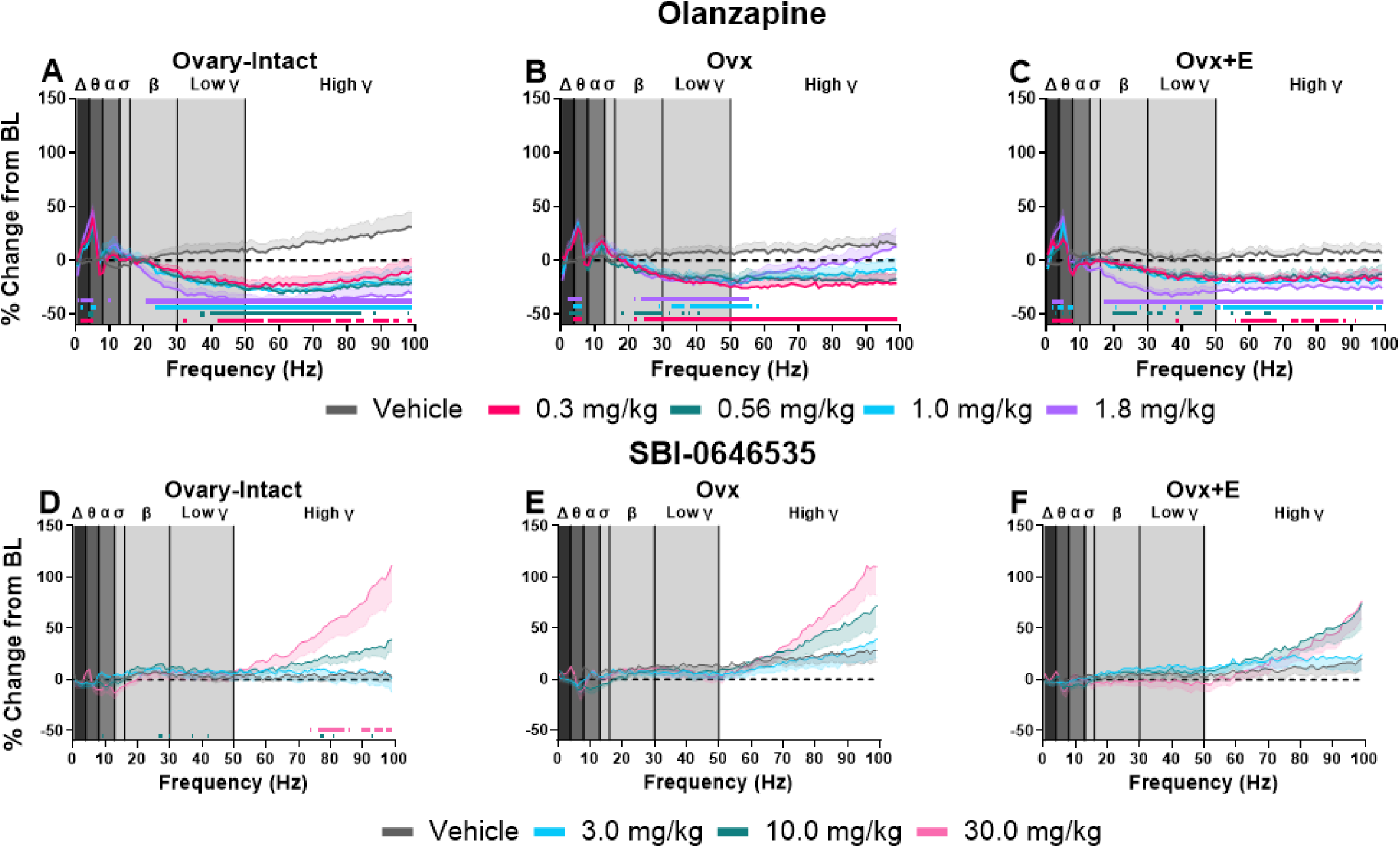
SBI-0646535 has a unique spectral profile compared to OLZ in all groups. All data are shown as a group mean (±SEM). Full spectrum data are presented in 1 Hz bins expressed as the average percent change from baseline during the 30-150 minute post-dosing period. Gray vertical bars represent frequency bands (delta, Δ 0.5–4 Hz; theta, θ 4–8 Hz; alpha, α 8–13 Hz; sigma, σ 13–15 Hz; beta, β 13–30 Hz; low gamma, γ 30–50 Hz; high gamma, γ 50–100 Hz). All tested OLZ doses were examined within each individual group: O-I (n= 9) (**A**), Ovx (n= 9) (**B**), and Ovx+E rats (n= 9) (**C**). All tested SBI-0646535 doses were also examined within each group: O-I (n = 8) (**D**), Ovx (n= 7-9) (**E**), and Ovx+E (n= 7-9) (**F**). p< 0.05; horizontal colored lines matching the color of respective doses represent frequencies at which OLZ- or SBI-0646535-treated groups were significantly different from vehicle-treated groups.

### SBI-0646535, but not OLZ, reduced gamma power equally across groups

Next, studies aimed to determine how efficacious these therapeutics may be to target NMDAR hypofunction associated with schizophrenia, especially considering the potential variability between populations/groups. Using qEEG as a functional biomarker of cortical activity, we evaluated efficacy of OLZ and SBI-0646535 to reduce MK-801-induced elevations in gamma power. In line with previous clinical reports of diminished antipsychotic response in postmenopausal patients, our findings directly support the hypothesis that OLZ would be less effective in attenuating MK-801-induced elevations in Ovx rats compared to other groups. First, OLZ significantly decreased MK-801-induced elevations in gamma power in all groups over time. Interestingly, while all tested doses were effective in O-I and Ovx+E rats, Ovx rats showed no significant reductions at the lowest dose. Further, there was a lower duration of effects at the 1.0 and 1.8 mg/kg dose in Ovx rats (>2 hours) compared to O-I (∼3 hours) and Ovx+E rats (5 hours). Together, these data suggest a need for higher antipsychotic doses in E2-deprived populations to elicit similar effects (**Fig. 5A-C**, for statistics see **Table 1**). Direct group comparisons in the high gamma power band further established this relationship, where OLZ significantly decreased MK-801-induced elevations in high gamma power across all tested doses in O-I and Ovx+E rats, but not Ovx rats (**Fig. 5D**).

**Figure 5.**
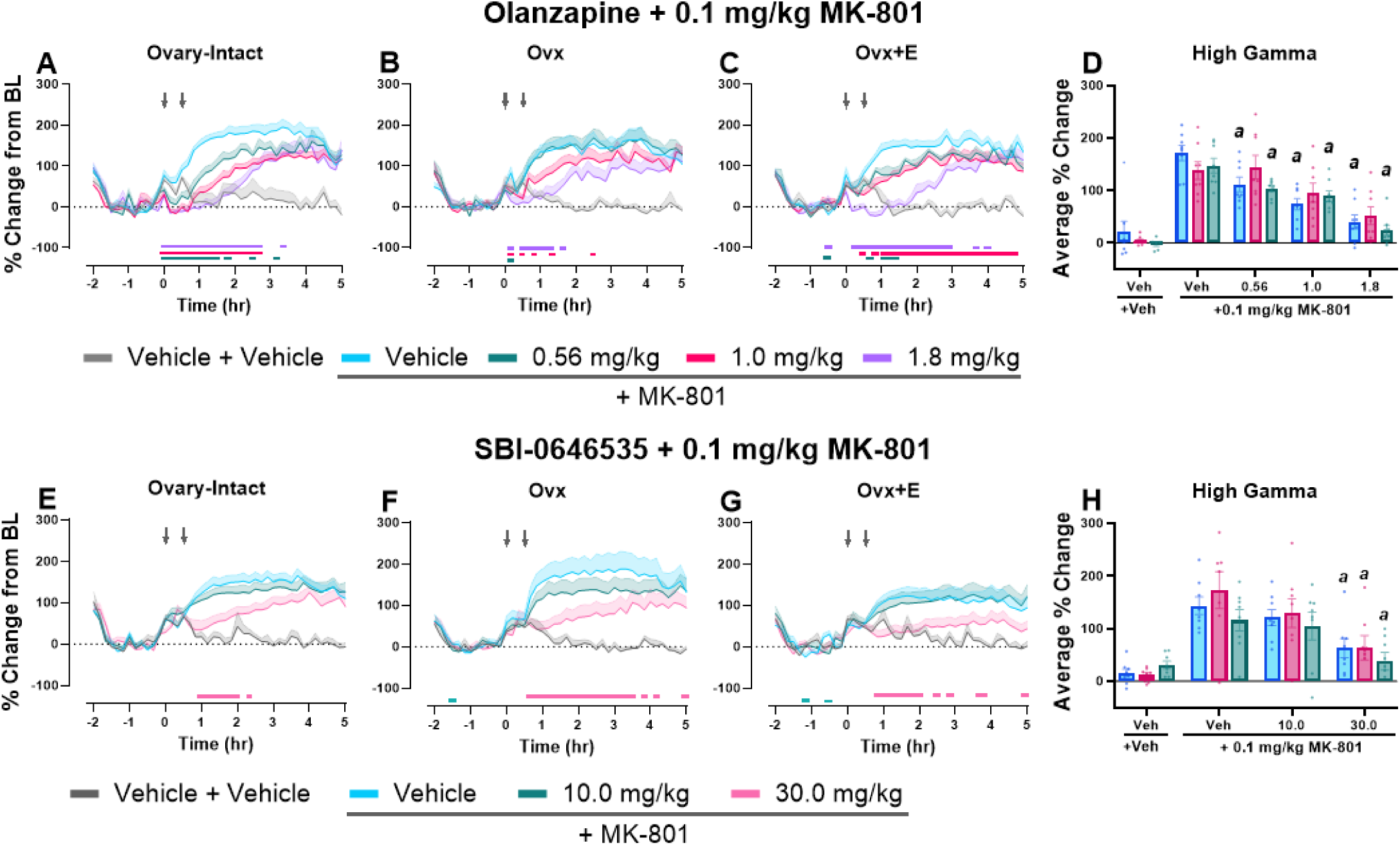
SBI-0646535, but not OLZ, was equally effective in mitigating MK-801-induced functional changes in Ovx rats. All data shown are group means ± SEM. The percent change from baseline in 10-minute bins across the 7-hour recording period was evaluated in O-I (**A,E**), Ovx (**B,F**), and Ovx+E rats (**C,G**). OLZ and SBI-0646535 were administered at time point 0 followed by vehicle or MK-801 30 minutes later (denoted by arrows). On the x-axis,-2 corresponds to ZT 0 and 5 corresponds to ZT 7. Then, direct group comparisons were evaluated as the average % change within the high gamma power band in the 60-180 min post-dosing period (**D**, main effect of dose: F_2.723, 57.19_= 44.29, p<0.0001; **H**, main effect of dose: F_1.753, 35.94_ = 32.96; p<0.0001). p<0.05; in timecourse graphs (**A-C** and **E-G**), horizontal colored lines matching the respective dose color represent the 10 min bins at which OLZ- or SBI-0646535-treated groups were significantly different from groups treated with vehicle + 0.1 mg/kg MK-801. In **D,H**, *a*, compared to the group’s respective vehicle + 0.1 mg/kg MK-801 condition. n=8 O-I, 7-8 Ovx, and 8 Ovx+E rats.

Despite observable differences between OLZ and SBI-0646535 on the spectral profile alone, SBI- 0646535 also significantly reduced MK-801-induced elevations in gamma power in all groups over time (for statistics, see **Table 2**). Significant reductions were observed at the 30 mg/kg dose in all groups (**Fig. 5E-H**). Unlike OLZ, both Ovx and Ovx+E rats showed significant reductions across the full 5 hour post-dosing period (**Fig. 5E-G**). Direct group comparisons in the high gamma power band revealed 30 mg/kg SBI-0646535 equally decreased MK-801-induced elevations in gamma power in the 60-180 min post-dosing period following (**Fig. 5H**). Thus, results support mGlu_2/3_ PAMs as a therapeutic approach effective regardless of hormonal status.

### Effects of MK-801 on locomotor activity and changes following administration of SBI-0646535 and OLZ were similar across groups

NMDAR antagonist-induced hyperlocomotion is frequently used as a preclinical behavioral readout for multiple symptoms of schizophrenia, and reductions in MK-801-induced increases in locomotor activity is commonly used to predict antipsychotic-like efficacy [20]. However, this paradigm often lacks translation as several compounds that reduce hyperlocomotor activity have demonstrated limited efficacy in clinical trials.

Thus, we primarily focused on functional readouts as a more translational approach and hypothesized that hyperlocomotion would be less sensitive to E2-associated variability between groups. Indeed, in our homecage activity assessment, MK-801 dose-dependently increased locomotor activity in all groups, but, unlike gamma power, there were no group differences (**Fig. 6A**). Interestingly, PEAQX and CP-101,606 had no effect on locomotor activity (**Fig. S13**)

**Figure 6.**
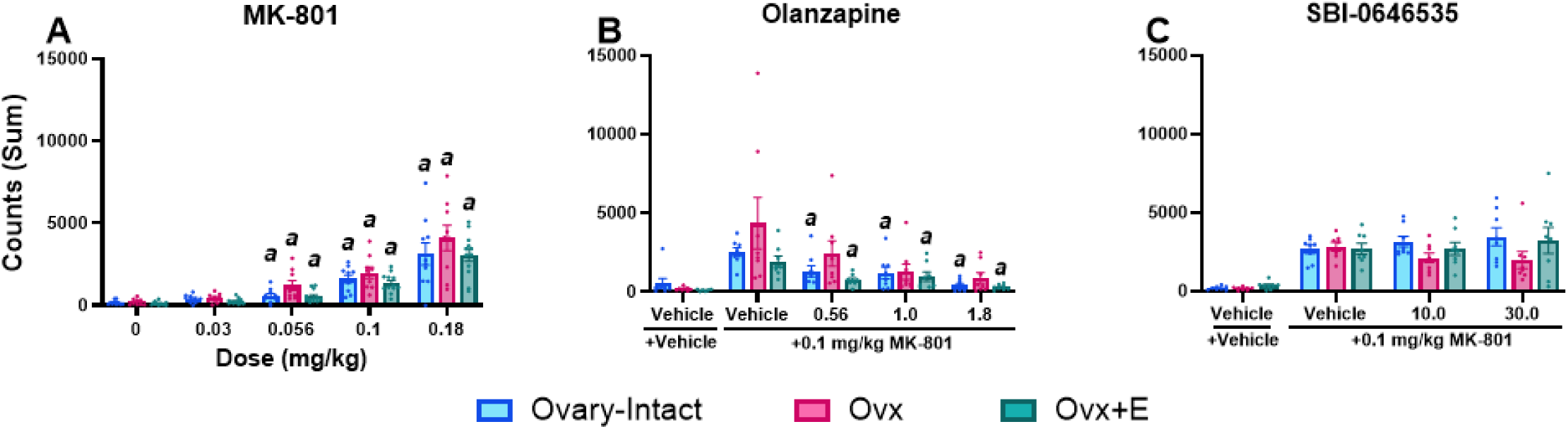
OLZ, but not SBI-0646535, reduced MK-801-induced increases in locomotor activity. For direct group comparisons, each individual’s locomotor activity counts were summed in the 30–90 min post-dosing period for MK-801 (main effect of dose: F_1.422, 39.46_= 67.00; p<0.0001) (**A**) or 60-180 min post dosing period for OLZ (main effect of dose: F_1.943, 39.82_= 11.79, p=0.0001) (**B)** or SBI-0636535 (**C**) combinations with MK-801 and graphed as a group mean ± SEM. *p* < 0.05; *a*, compared to group’s respective vehicle condition. For MK-801, n= 9-10 O-I, 8-9 Ovx, and 12 Ovx+E rats. For OLZ and SBI-0646535 combinations, n=8 O-I, 7-8 Ovx, and 8 Ovx+E rats.

In line with many other preclinical studies [20], we implemented hyperlocomotor activity as a screen to evaluate antipsychotic-like efficacy of OLZ and SBI-0646535 where we hypothesized both OLZ and SBI- 0646535 would reduce locomotor activity in all groups. As expected, OLZ did reduce MK-801-induced increases in locomotor activity, with significant reductions in O-I and Ovx+E rats, but not Ovx rats, at all tested doses (**Fig. 6B**); OLZ also reduced locomotor activity when administered alone in Ovx and Ovx +E rats (**Fig. S13**). Interestingly, in contrast to our hypothesis, SBI-0646535 had no effect on hyperlocomotor activity in any group/dose (**Fig. 6C**). Collectively, these results find a disconnect between locomotor and functional measures and suggest gamma power may be a more translational readout sensitive to differences in E2 status compared to classic locomotor assays.

## Discussion

Clinical evidence supports E2 deprivation as a risk factor for schizophrenia in females [5–7, 9–11, 13], yet the underlying mechanisms have not been extensively investigated. Findings from present studies show that Ovx alters NMDAR expression and function when probed by an NMDAR antagonist, and this effect is mitigated following E2 administration suggesting a primary role of E2. Notably, Ovx rats were more sensitive to MK-801-induced cognitive and functional disruptions that are, in part, a result of altered GluN2A expression.

These alterations may contribute to NMDAR hypofunction and could underlie increased symptom severity and diminished therapeutic response in patients with schizophrenia. Moreover, antipsychotic-like activity of OLZ was blunted in Ovx rats, whereas SBI-0646535 was effective in all conditions, suggesting mGlu_2/3_ PAMs may be a favorable therapeutic avenue in E2-deprived individuals.

First, we demonstrated that Ovx rats without E2 treatment were more sensitive to MK-801-induced disruptions on an attention task. Despite comparable dose-dependent increases in omissions in all groups, MK-801 reduced overall accuracy in O-I and Ovx rats, but not in Ovx+E rats on the 5CSRTT (**Fig. 1**). In fact, Ovx rats demonstrated the highest degree of attentional impairments, with dose-dependent decreases in accuracy spanning the 3 longer stimulus durations (**Fig. 1C-E**). Comparable to previous reports in male rats [48, 69], MK-801 dose-dependently reduced accuracy on the PAL task in O-I and Ovx rats but, interestingly, did not impact accuracy in Ovx+E rats except at the 0.18 mg/kg dose which also drastically impacted trial completion (**Fig. 2**). Importantly, MK-801-induced homecage hyperlocomotor activity was not different between groups (**Fig. 6**), suggesting that altering E2 status selectively impacted cognitive performance. Overall, these results align with prior preclinical studies that show, under the appropriate dosing conditions, E2 administration to Ovx animals can improve cognitive impairments associated with Ovx alone [36, 50, 74–76]. However, our studies are the first to examine NMDAR antagonist-induced cognitive disruptions on these translational touchscreen tasks, extending prior work to establish the relationship between NMDAR function and E2 status.

Next, we showed that Ovx rats without E2 treatment were more sensitive to MK-801-induced elevations in gamma power, a cortical biomarker of E/I balance widely disrupted in patients with schizophrenia. As expected, MK-801 dose-dependently increased high frequency gamma power in all groups. However, at the highest dose administered (0.18 mg/kg), Ovx rats showed a blunted response to MK-801 compared to the Ovx+E rats, regarding both peak effect (**Fig. 2B, C**) and timecourse (**Fig 2E**). This inverted-U response may be explained by shifts in NMDAR blockade within cortical disinhibitory circuits. Gamma band oscillations are hypothesized to be generated by the reciprocal activity of PV interneurons and glutamatergic pyramidal neurons [26, 29, 77]. At low to moderate doses, NMDAR antagonists preferentially block NMDARs on PV interneurons [77–79], resulting in disinhibition of pyramidal cells and increased excitation as demonstrated at the 0.03-0.1 mg/kg doses in present studies. In further support of the role of NMDARs on PV interneurons, mice with a genetic knockdown of NMDARs on PV cells show elevated resting-state gamma power [28]. The contrasting blunted elevations associated with higher doses of NMDAR antagonists are hypothesized to be driven by increased inhibition of NMDARs on pyramidal neurons. This dual blockade counteracts disinhibitory effects, reducing the associated E/I ratio [26]. Thus, results indicate a leftward-shift in Ovx animals at the highest tested dose (0.18 mg/kg).

Observed changes in E/I balance following Ovx suggest altered NMDAR expression and/or function. Indeed, findings confirmed that Ovx resulted in lower GluN2A and GluN2B NMDAR synaptic expression in the PFC that was rescues with E2 treatment. Furthermore, qEEG studies revealed a heightened sensitivity of GluN2A, but not GluN2B inhibition, in Ovx rats. Taken together, these findings suggest E2 depletion exacerbates response to NMDAR antagonism, and this is in part regulated by altered GluN2A expression/function. These align with postmortem studies in patients with schizophrenia that reported reduced GluN2A mRNA expression on cortical PV interneurons [32]. Given that GluN2A is the dominant subunit on PV interneurons [80], future studies are needed to directly examine NMDAR subunit expression and function specifically on PV interneurons.

Notably, PEAQX and MK-801 yielded distinct effects on gamma power. Unlike MK-801, group differences in response to PEAQX were associated with overall magnitude of percent change rather than a leftward-shift in the dose-response curve. It is unknown if PEAQX produces a biphasic dose response on gamma power like MK-801; higher doses were not administered based on prior reports of toxicity [52].

Additionally, effects of PEAQX were most prevalent in the low, not high, gamma frequency range, yet the inverse was true for MK-801. The molecular mechanisms regulating low versus high gamma oscillations are not well understood, but this distinction suggests GluN2A strongly mediates low gamma oscillatory activity. Both low and high gamma power abnormalities have been reported in neuropsychiatric disorders including schizophrenia and both are linked to associated cognitive dysfunction [72, 81]. Nevertheless, these biochemical and functional results support clear interactions between E2, NMDAR function, and E/I balance.

Finally, present results support further examination of mGlu_2/3_ PAMs as a novel target with superior antipsychotic-like activity for E2-deprived individuals. In all groups, OLZ shifted relative power from high to low frequencies (**Fig. 4A-C**), consistent with a sedative profile and a commonly reported side effect of antipsychotic medications [82–85]. In contrast, SBI-0646535 did not alter low frequency power demonstrating a potential for lesser sedative effects (**Fig. 4D-F**). Interestingly, SBI-0646535 increased high frequency gamma power only at and above 70 Hz. In contrast to MK-801, doses of SBI-0646535 that elevated this frequency range did not induce hyperlocomotor activity (**Fig. 6**) nor disrupt working memory (**Fig. S2**). Thus, these effects may be promising as moderate elevations in gamma power can be beneficial and increases in both low and high gamma oscillations are critical in functions including sensory processing and working memory [72, 86–89].

However, aberrant elevations in gamma power are associated with symptoms of schizophrenia and have been used as a targetable pharmacodynamic biomarker to predict treatment efficacy [90–92]. Consistent with previous literature evaluating antipsychotic medications in male rats [57, 82, 93, 94], OLZ attenuated MK- 801-induced increases in gamma power at a dose that produced comparable elevations in all groups.

However, Ovx rats required higher doses of OLZ to elicit a similar magnitude of effects as those observed in Ovx+E and O-I. In contrast, SBI-0646535 equally attenuated MK-801-induced elevations in gamma power in all groups. Thus, E2-depletion may have a causal role in the reduced sensitivity of antipsychotic medications in postmenopausal female patients [13], whereas E2-depletion did not affect sensitivity to mGlu_2/3_ PAMs. These results also strengthen support for the use of gamma power as a translational biomarker over historical assessments of hyperlocomotion to aid therapeutic development. Regardless of E2 status, MK-801 induced similar increases in locomotor activity within the homecage, yet elevations in gamma power revealed group differences. Consistent with prior literature, OLZ attenuated MK-801-induced hyperlocomotor activity, yet the mGlu_2/3_ PAM did not, revealing a disconnect between effects on locomotion and gamma power.

Collectively, our data support the hypothesis that E2 depletion may serve as a risk factor for schizophrenia by impacting NMDAR activity; these effects are sufficiently mediated by altered GluN2A expression and function. Furthermore, exogenous E2 administration in most instances restored effects in Ovx rats, aligning with findings from O-I rats. While results do support a strong role of E2, future studies are needed to interrogate the effects of progesterone. Few clinical studies have evaluated relationships between progesterone and schizophrenia, with existing studies yielding mixed results; some studies have reported increased symptom severity in premenopausal patients during the low progesterone phase of their menstrual cycle [95, 96]. Additionally, while Ovx was used to systematically evaluate complete ovarian hormone loss and subsequent contributions of E2 mapped onto this ovarian hormone “blank-slate”, a valuable future direction is to expand studies using the 4-vinylcyclohexene diepoxide (VCD) rodent model. The VCD model is highly translational and yields perimenopausal-like hormonal fluctuations [38]. Moreover, while hormone therapy has been effective in improving overall symptom outcome in patients with schizophrenia and present studies support a beneficial role of E2, there are risks associated with long-term E2 treatment, and if the uterus is present a progestogen must be dually administered. Together, this may outweigh the associated therapeutic benefits [13, 97, 98]; few clinical studies have evaluated this in schizophrenia, with one study finding E2 combined with medroxyprogesterone acetate (MPA) improved negative and cognitive symptoms in female patients [99] and a separate study reporting no benefits [100]. Thus, novel, non-hormonal therapeutic approaches may be better in postmenopausal populations.

Overall, there is great need to decipher a multi-systems risk to benefit personalized medicine framework, as well as develop more tolerable and effective antipsychotic medications for individuals with schizophrenia, especially those with low E2 levels. Present studies support further development of mGlu_2/3_ PAMs as a novel antipsychotic mechanism whereby efficacy, to date, is not impacted by E2 status and may be a safer alternative to adjunctive estrogen treatment. As precision medicine has become an increasing priority in drug development, acknowledging how factors such as hormone status interact to influence course of illness is a critical next step to develop treatments and enhance quality of life as aging ensues.

## Author Contributions

**Kimberly Holter**: conceptualization, formal analysis, investigation, methodology, visualization, writing – original draft and reviewing & editing. **McKenna Klausner and Mary Hunter Hite**: investigation and formal analysis. **Samuel Barth**: investigation and formal analysis. **Bethany Pierce**: investigation. **Alexandria Iannucci:** investigation. **Douglas Sheffler:** Resources. **Nicholas Cosford:** Resources. **Heather Bimonte-Nelson:** conceptualization, methodology. **Kimberly Raab-Graham:** conceptualization, methodology, formal analysis **Robert Gould**: conceptualization, methodology, formal analysis, visualization, writing – review and editing, funding acquisition.

## Acknowledgements

We wish to thank Drs. Justin Strickland and Paul Sands for their assistance in writing custom R and MATLAB scripts for data analysis. Further, we wish to thank Owen Ghaphery, Ashlyn Stone, and Elizabeth Bedingham for their efforts involved in the manual sleep scoring for qEEG studies as well as Dr. Liliya Yamaleyeva for providing training on ovariectomization procedures.

