## Supplemental Material for "17β-estradiol status alters NMDAR function and antipsychotic-like activity in female rats"

**Supplemental Methods:**

*17β-Estradiol administration*

Silastic capsules were made as previously described [1,2]. 25% E2 (17β-Estradiol; Sigma-Aldrich) and 75% cholesterol (Sigma-Aldrich) were dissolved in 200-proof ethanol. Following ethanol evaporation the remaining powder was packed into 5 mm of Silastic tubing (VWR, Randor, PA; Dow Corning; inner diameter: 0.058″ outer diameter: 0.077″ wall thickness: 0.009″). Capsules were secured tightly on both ends using wooden dowels (1-2mm) and silicone glue and checked for leaks once dry. E2-containing capsules were soaked in sterile saline for 24 hours prior to implantation. For ovary-intact and Ovx groups, capsules remained empty as no significant effects have been reported using cholesterol alone as a vehicle condition [1,3]. Neuroendocrine status for successful Ovx (diestrus-like) and E2 replacement (estrus-like) was confirmed using vaginal cytology.

*Vaginal cytology*

Vaginal lavages were conducted to assess estrous cycle to (1) validate surgical procedures (Ovx: diestrus-like; Ovx+E: estrus-like) and (2) prior to all dosing studies for ovary-intact rats. Rats were gently picked up by the base of the tail and the tip of a glass Pasteur pipette was inserted ~1-2 mm into the vaginal canal. Around 0.1 ml of room temperature sterile saline was flushed in and out of the cavity 3 times, and the fluid from the lavage was then placed on a microscope slide. Cells were viewed under a light microscope and classified as metestrus-, diestrus-, proestrus-, or estrus-like based on characteristic patterns previously described by Koebele and Bimonte-Nelson 2016.

*Touchscreen training*

5-choice Serial Reaction Time Task (5CSRTT)

Rats (Ovx+E n=8, Ovx n=13, Ovary-Intact n=10) were trained 6-7 days per week during the first half of their light cycle to touch stimuli presented on the touchscreen in operant chambers (Lafayette Instruments, Lafayette, IN) for a 45 mg chocolate sucrose pellet reward (Bio-Serv). Training involved Initial Touch Training (ITT) and Must Touch Training (MTT) followed by rat touch basic training. In ITT, an illuminated square was presented in one of five windows spaced horizontally across the computer touchscreen. Touching the square resulted in 3 sucrose pellets presented in the illuminated magazine. No response after 30 seconds resulted in stimulus removal and presentation of 1 sucrose pellet into the magazine located on the opposite side of the chamber. A nose-poke in the magazine to consume the reward resulted in the light turning off, a “click” noise, and initiation of the next trial. In MTT, rats were required to touch the illuminated square to receive a 1 sucrose pellet reward. A minimum of 30 trials had to be completed to move to the next stage.

Rats then underwent basic training which consisted of 8 phases. In each phase, rats were presented an illuminated square in one of five windows in a pseudorandom order. Touching the location where the square was illuminated within the limited hold period resulted in illumination of the receptacle light, a 1-second tone, and reward delivery. An incorrect response (touching the incorrect location) within the limited hold period or omission (no response after the limited hold period time had elapsed) resulted in the illumination of the house light for 5 seconds and a 20-second time out period. The duration the stimulus was illuminated decreased following advancement to each phase (60 seconds in session 1 to 1 second in session 8). The limited hold period, time following stimulus presentation in which the rat was allowed to respond, also decreased (60 seconds in session 1 to 5 seconds in session 8). Rats had to perform ≥ 80% accuracy and ≤20% omissions to progress to the next phase. After successful completion of all 8 phases, rats advanced to the variable task.

Paired Associates Learning (PAL) Task

Rats (Ovx n=10, Ovx+E n=13, and ovary-intact n=10) underwent touchscreen training as previously described [4,5]. Following initial and must touch training as above (ITT and MTT), rats had to learn to initiate each trial by nosepoking in the magazine opposite the touchscreen (MIT) and successfully completing 30 trails for 2 sessions. Lastly, rats had to learn to track and respond on an illuminated stimulus as opposed to a non-illuminated square at >80% accuracy for 2 sessions (punish incorrect training; PIT), Following successful completion of all training phases rats advanced to the PAL task.

**Table S1.** Statistics describing the outcomes of MK-801 on relative power spectral distribution across groups.

| **Full Spectrum Statistics** | | | | | | | |
|---|---|---|---|---|---|---|---|
| **Figure** | **Source of Variation** | **DF** | **F** | **p** | ***** | ***Post hoc* results** | **Significant Frequencies (Hz)** |
| **S3A MK-801 Ovary-Intact** | Group | 2.50, 22.52 | 13.00 | <0.0001 | **** | Vehicle vs. 0.03 mg/kg<br>Vehicle vs. 0.056 mg/kg<br>Vehicle vs. 0.1 mg/kg<br>Vehicle vs. 0.18 mg/kg | 6, 7, 10-19, 30-92, 96<br>5-7, 10-19, 33-95<br>0.5-9, 11-17, 19, 28-97<br>6-10, 13-22, 38-99 |
| | Frequency | 2.54, 22.83 | 124.7 | <0.0001 | **** | | |
| | Interaction | 4.11, 33.88 | 10.52 | <0.0001 | **** | | |
| **S3B MK-801 Ovx** | Group | 2.84, 22.74 | 8.582 | 0.0006 | *** | Vehicle vs. 0.03 mg/kg<br>Vehicle vs. 0.056 mg/kg<br>Vehicle vs. 0.1 mg/kg<br>Vehicle vs. 0.18 mg/kg | 4-6, 11-18, 31, 32, 34-94, 96-98<br>5, 11-16, 18, 19, 33-99<br>2, 3, 5-9, 12-22, 31-99<br>2, 6-10, 14-25, 27, 28, 43-99 |
| | Frequency | 2.03, 16.24 | 23.84 | <0.0001 | **** | | |
| | Interaction | 4.58, 36.65 | 8.086 | <0.0001 | **** | | |
| **S3C MK-801 Ovx+E** | Group | 2.42, 26.65 | 18.84 | <0.0001 | **** | Vehicle vs. 0.03 mg/kg<br>Vehicle vs. 0.056 mg/kg<br>Vehicle vs. 0.1 mg/kg<br>Vehicle vs. 0.18 mg/kg | 4-6, 10-14, 16, 18-21, 32, 33, 35-93, 95<br>1-6, 8, 10-19, 28-99<br>0.5-6, 8, 9, 12-22, 28-99<br>1-10, 13-21, 30, 31, 35-99 |
| | Frequency | 3.52, 38.67 | 71.16 | <0.0001 | **** | | |
| | Interaction | 4.67, 51.38 | 12.27 | <0.0001 | **** | | |
| *p<0.05; ** p<0.01; *** p<0.001; ****p<0.0001 | | | | | | | |

**Table S2.** Statistics for NMDA receptor antagonist timecourse graphs (MK-801, CP-101,606, and PEAQX).

| Timecourse Statistics | | | | | | | |
|---|---|---|---|---|---|---|---|
| **Figure** | **Source of Variation** | **DF** | **F** | **p** | ***** | ***Post hoc* results** | **Significant Timepoints (10-minute bin)** |
| **MK-801** | | | | | | | |
| **Ovary-Intact** | | | | | | | |
| **S5A** Delta | Dose | 2.49, 22.38 | 7.670 | 0.0017 | ** | Vehicle vs 0.03 mg/kg | 30, 40 |
| | Time | 5.72, 51.50 | 7.266 | <0.0001 | **** | Vehicle vs 0.1 mg/kg | 20-150, 170, 260 |
| | Interaction | 6.61, 53.98 | 2.837 | 0.0151 | * | Vehicle vs 0.18 mg/kg | 30-60, 110-300 |
| **S5D** Theta | Dose | 2.89, 25.82 | 98.74 | <0.0001 | **** | Vehicle vs 0.03 mg/kg | 40-60, 100 |
| | Time | 3.92, 35.29 | 39.46 | <0.0001 | **** | Vehicle vs 0.056 mg/kg | 30, 50-180, 250 |
| | Interaction | 6.98, 57.02 | 13.19 | <0.0001 | **** | Vehicle vs 0.1 mg/kg | 10-300 |
| | | | | | | Vehicle vs 0.18 mg/kg | 10-300 |
| **S5G** Alpha | Dose | 2.20, 19.84 | 14.10 | 0.0001 | *** | Vehicle vs 0.03 mg/kg | -40, 120, 130 |
| | Time | 4.86, 43.69 | 5.293 | 0.0008 | *** | Vehicle vs 0.056 mg/kg | -30, 130, 150, 160, 250-270 |
| | Interaction | 6.98, 57.07 | 3.586 | 0.0029 | ** | Vehicle vs 0.1 mg/kg | 50, 110, 130, 160 |
| | | | | | | Vehicle vs 0.18 mg/kg | 10, 30-60, 100, 110, 130, 140, 160, 180-220 |
| **S5J** Sigma | Dose | 2.13, 19.20 | 30.31 | <0.0001 | **** | Vehicle vs 0.03 mg/kg | 30-60, 80, 90, 110, 130 |
| | Time | 5.31, 47.81 | 13.72 | <0.0001 | **** | Vehicle vs 0.056 mg/kg | 10-30, 50-170, 230, 250 |
| | Interaction | 7.39, 60.36 | 5.047 | 0.0001 | *** | Vehicle vs 0.1 mg/kg | 10-300 |
| | | | | | | Vehicle vs 0.18 mg/kg | 0, 10, 60, 70, 80-100, 130, 150-170, 210-300 |
| **S5M** Beta | Dose | 2.53, 22.73 | 9.798 | 0.0004 | *** | Vehicle vs 0.03 mg/kg | 20, 50, 60, 160, 230, 290 |
| | Time | 3.75, 33.72 | 4.695 | 0.0047 | ** | Vehicle vs 0.056 mg/kg | 20, 30, 50-100, 130, 140, 160-180, 270 |
| | Interaction | 5.23, 42.71 | 2.088 | 0.0829 | ns | Vehicle vs 0.1 mg/kg | 50, 80, 90, 130, 160-180, 220-300 |
| | | | | | | Vehicle vs 0.18 mg/kg | 10-100, 120-140, 160, 170, 230, 240, 270, 290 |
| **S5P** Low Gamma | Dose | 2.15, 19.32 | 27.17 | <0.0001 | **** | Vehicle vs 0.03 mg/kg | 30-50, 70-110, 130 |
| | Time | 4.81, 43.31 | 22.41 | <0.0001 | **** | Vehicle vs 0.056 mg/kg | -60, 20-40, 60-200 |
| | Interaction | 6.53, 53.40 | 6.533 | <0.0001 | **** | Vehicle vs 0.1 mg/kg | 10-300 |
| | | | | | | Vehicle vs 0.18 mg/kg | 0, 10, 30-300 |
| **Ovx** | | | | | | | |
| **S5B** Delta | Dose | 1.91, 15.25 | 2.375 | 0.1282 | ns | | |
| | Time | 2.99, 23.89 | 4.387 | 0.0136 | * | | |
| | Interaction | 5.91, 45.17 | 1.913 | 0.1002 | ns | | |
| **S5E** Theta | Dose | 1.99, 15.88 | 4.035 | 0.0386 | * | Vehicle vs 0.03 mg/kg | 70, 100-120, 150, 190, 200, 220 |
| | Time | 3.03, 24.21 | 7.622 | 0.0009 | *** | Vehicle vs 0.056 mg/kg | 70, 120, 150, 180-200, 220, 230, 250, 290 |
| | Interaction | 5.08, 38.98 | 2.408 | 0.0529 | ns | Vehicle vs 0.1 mg/kg | 40-80, 100-300 |
| | | | | | | Vehicle vs 0.18 mg/kg | -50, 200, 220, 230, 250, 280-300 |
| **S5H** Alpha | Dose | 2.58, 20.67 | 10.09 | 0.0004 | *** | Vehicle vs 0.03 mg/kg | -110, 190, 200, 220-250 |
| | Time | 3.34, 26.71 | 6.198 | 0.0019 | ** | Vehicle vs 0.056 mg/kg | 190 |
| | Interaction | 5.25, 40.29 | 4.685 | 0.0016 | ** | Vehicle vs 0.1 mg/kg | -110, 50 |
| | | | | | | Vehicle vs 0.18 mg/kg | 30, 50, 70-170 |
| **S5K** Sigma | Dose | 2.71, 21.65 | 22.43 | <0.0001 | **** | Vehicle vs 0.03 mg/kg | 30-60, 80, 90, 130-150, 170-190, 210, 230-250 |
| | Time | 3.74, 29.88 | 10.14 | <0.0001 | **** | Vehicle vs 0.056 mg/kg | 20, 40, 60, 110-260, 290 |
| | Interaction | 5.78, 44.37 | 4.24 | 0.0021 | ** | Vehicle vs 0.1 mg/kg | 10-90, 110-300 |
| | | | | | | Vehicle vs 0.18 mg/kg | 60-90, 110-260, 280, 290 |
| **S5N** Beta | Dose | 3.09, 24.69 | 16.41 | <0.0001 | **** | Vehicle vs 0.03 mg/kg | 90, 130, 140, 200, 260 |
| | Time | 3.59, 28.73 | 8.735 | 0.0001 | *** | Vehicle vs 0.056 mg/kg | 110, 130-170, 230, 280 |
| | Interaction | 5.30, 40.53 | 2.840 | 0.0252 | * | Vehicle vs 0.1 mg/kg | 10, 20, 60-90, 110-180, 200, 220-290 |
| | | | | | | Vehicle vs 0.18 mg/kg | 20-200, 220-290 |
| **S5Q** Low Gamma | Dose | 2.53, 20.22 | 12.13 | 0.0002 | *** | Vehicle vs 0.03 mg/kg | 50, 100, 210, 220, 240 |
| | Time | 3.96, 31.68 | 11.50 | <0.0001 | **** | Vehicle vs 0.056 mg/kg | 50-70, 90-260, 280, 290 |
| | Interaction | 5.98, 45.90 | 4.735 | 0.0008 | *** | Vehicle vs 0.1 mg/kg | 10-300 |
| | | | | | | Vehicle vs 0.18 mg/kg | 190-260, 280-300 |
| **Ovx+E** | | | | | | | |
| **S5C** Delta | Dose | 3.21, 35.32 | 22.92 | <0.0001 | **** | Vehicle vs 0.03 mg/kg | 230 |
| | Time | 7.17, 78.87 | 11.16 | <0.0001 | **** | Vehicle vs 0.056 mg/kg | 50, 60 |
| | Interaction | 8.26, 90.27 | 4.47 | 0.0001 | *** | Vehicle vs 0.1 mg/kg | 30-210, 240 |
| | | | | | | Vehicle vs 0.18 mg/kg | 30-60, 110-300 |
| **S5F** Theta | Dose | 1.98, 21.73 | 31.32 | <0.0001 | **** | Vehicle vs 0.03 mg/kg | 40, 50, 80, 90, 110-130, 150, 190, 200, 220 |
| | Time | 5.64, 62.00 | 26.90 | <0.0001 | **** | Vehicle vs 0.056 mg/kg | 30-50, 70-260, 280-300 |
| | Interaction | 7.49, 81.93 | 7.349 | <0.0001 | **** | Vehicle vs 0.1 mg/kg | -70, 10-300 |
| | | | | | | Vehicle vs 0.18 mg/kg | 30-300 |
| **S5I** Alpha | Dose | 2.25, 24.69 | 23.90 | <0.0001 | **** | Vehicle vs 0.03 mg/kg | 90, 210 |
| | Time | 4.15, 45.65 | 15.02 | <0.0001 | **** | Vehicle vs 0.056 mg/kg | -60, -30, 210, 220, 240, 260-280 |

| | | | | | | | |
|---|---|---|---|---|---|---|---|
| | Interaction | 6.02, 65.80 | 9.072 | <0.0001 | **** | Vehicle vs 0.1 mg/kg<br>Vehicle vs 0.18 mg/kg | 20-120, 140, 160-190<br>-30, 10-300 |
| **S5L**<br>Sigma | Dose | 2.47, 27.19 | 20.01 | <0.0001 | **** | Vehicle vs 0.03 mg/kg<br>Vehicle vs 0.056 mg/kg<br>Vehicle vs 0.1 mg/kg<br>Vehicle vs 0.18 mg/kg | 60, 70, 90-110, 130, 140, 180-210, 230, 240, 260, 270<br>40-110, 130-290<br>20-300<br>10, 60, 100, 150, 180-300 |
| | Time | 5.57, 61.23 | 11.74 | <0.0001 | **** | | |
| | Interaction | 8.46, 92.55 | 5.412 | <0.0001 | **** | | |
| **S5O**<br>Beta | Dose | 2.65, 29.18 | 11.04 | <0.0001 | **** | Vehicle vs 0.03 mg/kg<br>Vehicle vs 0.056 mg/kg<br>Vehicle vs 0.1 mg/kg<br>Vehicle vs 0.18 mg/kg | 90, 100, 130, 220, 230, 270, 290<br>90-140, 240, 250, 270<br>50, 80-140, 160-180, 200-300<br>50-290 |
| | Time | 6.27, 69.01 | 4.658 | 0.0004 | *** | | |
| | Interaction | 9.02, 98.57 | 2.849 | 0.005 | ** | | |
| **S5R**<br>Low Gamma | Dose | 1.53, 16.87 | 17.51 | <0.0001 | **** | Vehicle vs 0.03 mg/kg<br>Vehicle vs 0.056 mg/kg<br>Vehicle vs 0.1 mg/kg<br>Vehicle vs 0.18 mg/kg | 60, 70, 170<br>20-220, 240-290<br>10-300<br>10, 20, 40-70, 110, 130, 150, 170-300 |
| | Time | 6.52, 71.68 | 32.44 | 0.0002 | *** | | |
| | Interaction | 7.09, 77.56 | 6.87 | <0.0001 | **** | | |
| **PEAQX** | | | | | | | |
| **Ovary-Intact** | | | | | | | |
| **S9A**<br>Delta | Dose | 3.58, 25.04 | 0.6545 | 0.4701 | ns | | |
| | Time | 1.22, 8.504 | 1.326 | 0.2881 | ns | | |
| | Interaction | 5.12, 32.86 | 1.229 | 0.3176 | ns | | |
| **S9D**<br>Theta | Dose | 1.99, 13.95 | 11.51 | 0.0011 | ** | Vehicle vs 10.0 mg/kg<br>Vehicle vs 30.0 mg/kg | -60, 70-100<br>-80, 60-120, 140, 210 |
| | Time | 5.07, 35.47 | 1.874 | 0.1229 | ns | | |
| | Interaction | 5.80, 37.21 | 1.742 | 0.1406 | ns | | |
| **S9G**<br>Alpha | Dose | 1.93, 13.49 | 14.53 | 0.0005 | *** | Vehicle vs 10.0 mg/kg<br>Vehicle vs 30.0 mg/kg | 20, 30, 60, 70, 130, 140<br>10-40, 60-90, 140, 160, 180, 240 |
| | Time | 4.35, 30.43 | 2.924 | 0.0337 | * | | |
| | Interaction | 5.85, 37.54 | 1.951 | 0.0992 | ns | | |
| **S9J**<br>Sigma | Dose | 1.68, 11.78 | 11.02 | 0.0027 | ** | Vehicle vs 10.0 mg/kg<br>Vehicle vs 30.0 mg/kg | 60, 130, 150<br>20, 30, 50-90, 110, 120, 160, 250 |
| | Time | 5.03, 35.21 | 7.220 | <0.0001 | **** | | |
| | Interaction | 5.31, 34.06 | 1.618 | 0.1788 | ns | | |
| **S9M**<br>Beta | Dose | 1.63, 11.39 | 2.623 | 0.122 | ns | | |
| | Time | 4.78, 33.43 | 5.508 | 0.001 | *** | | |
| | Interaction | 5.30, 34.00 | 1.246 | 0.3093 | ns | | |
| **S9P**<br>Low Gamma | Dose | 1.29, 9.051 | 8.884 | 0.0118 | * | Vehicle vs 10.0 mg/kg<br>Vehicle vs 30.0 mg/kg | 30<br>20, 30, 50-80, 110, 120, 140, 160-180 |
| | Time | 4.84, 33.89 | 3.514 | 0.0121 | * | | |
| | Interaction | 4.98, 31.94 | 2.163 | 0.0834 | ns | | |
| **S9S**<br>High Gamma | Dose | 1.37, 9.611 | 9.176 | 0.0093 | ** | Vehicle vs 10.0 mg/kg<br>Vehicle vs 30.0 mg/kg | -80, 60<br>20, 30, 60-80, 120 |
| | Time | 4.31, 30.13 | 8.180 | 0.0001 | *** | | |
| | Interaction | 5.22, 33.48 | 2.244 | 0.0703 | ns | | |
| **Ovx** | | | | | | | |
| **S9B**<br>Delta | Dose | 1.27, 8.863 | 2.847 | 0.1224 | ns | | |
| | Time | 4.71, 32.93 | 4.174 | 0.0054 | ** | | |
| | Interaction | 4.86, 33.58 | 1.722 | 0.1581 | ns | | |
| **S9E**<br>Theta | Dose | 1.04, 7.274 | 25.68 | 0.0012 | ** | Vehicle vs 10.0 mg/kg<br>Vehicle vs 30.0 mg/kg | 40, 60, 90, 100, 150<br>-100, -50, 30-60, 80-100, 120-270, 290, 300 |
| | Time | 4.77, 33.36 | 7.416 | 0.0001 | *** | | |
| | Interaction | 6.09, 42.08 | 4.019 | 0.0027 | ** | | |
| **S9H**<br>Alpha | Dose | 1.94, 13.56 | 4.932 | 0.0254 | * | Vehicle vs 10.0 mg/kg<br>Vehicle vs 30.0 mg/kg | 100<br>90, 100, 160, 180, 190, 250-270, 300 |
| | Time | 4.39, 30.74 | 3.801 | 0.0108 | * | | |
| | Interaction | 5.60, 38.69 | 1.711 | 0.1486 | ns | | |
| **S9K**<br>Sigma | Dose | 1.94, 13.56 | 4.932 | 0.0254 | * | Vehicle vs 10.0 mg/kg<br>Vehicle vs 30.0 mg/kg | 100<br>90, 100, 160, 180, 190, 250-270, 300 |
| | Time | 4.39, 30.74 | 3.801 | 0.0108 | * | | |
| | Interaction | 5.60, 38.69 | 1.711 | 0.1486 | ns | | |
| **S9N**<br>Beta | Dose | 1.67, 11.67 | 0.8474 | 0.4337 | ns | | |
| | Time | 3.47, 24.30 | 4.511 | 0.0093 | ** | | |
| | Interaction | 4.51, 31.15 | 1.178 | 0.3412 | ns | | |
| **S9Q**<br>Low Gamma | Dose | 1.52, 10.67 | 8.103 | 0.0102 | * | Vehicle vs 10.0 mg/kg<br>Vehicle vs 30.0 mg/kg | 90, 100<br>80-110, 130-180, 250, 270, 300 |
| | Time | 4.78, 33.48 | 5.811 | 0.0007 | *** | | |
| | Interaction | 5.11, 35.25 | 3.339 | 0.0138 | * | | |

| | | | | | | | |
|---|---|---|---|---|---|---|---|
| **S9T** High Gamma | Dose | 1.46, 10.24 | 3.834 | 0.067 | ns | | |
| | Time | 4.79, 33.53 | 4.225 | 0.0048 | ** | | |
| | Interaction | 5.20, 35.97 | 2.173 | 0.0766 | ns | | |
| **Ovx+E** | | | | | | | |
| **S9C** Delta | Dose | 1.24, 8.663 | 3.656 | 0.0843 | ns | | |
| | Time | 4.85, 33.95 | 6.054 | 0.0005 | *** | | |
| | Interaction | 5.06, 34.96 | 1.515 | 0.2098 | ns | | |
| **S9F** Theta | Dose | 1.32, 9.255 | 11.74 | 0.0051 | ** | Vehicle vs 10.0 mg/kg<br>Vehicle vs 30.0 mg/kg | 200, 300<br>30-50, 100, 110, 160-190, 210, 220, 240, 270, 280 |
| | Time | 4.03, 28.18 | 3.580 | 0.0174 | * | | |
| | Interaction | 5.65, 39.06 | 1.645 | 0.1644 | Ns | | |
| **S9I** Alpha | Dose | 1.24, 8.699 | 18.19 | 0.0016 | ** | Vehicle vs 10.0 mg/kg<br>Vehicle vs 30.0 mg/kg | -50, 170<br>30, 40, 60-80, 120, 160, 170, 250 |
| | Time | 3.31, 23.18 | 7.721 | 0.0007 | *** | | |
| | Interaction | 5.61, 38.81 | 1.569 | 0.1858 | Ns | | |
| **S9L** Sigma | Dose | 1.88, 13.19 | 2.780 | 0.1005 | ns | | |
| | Time | 4.75, 33.22 | 6.520 | 0.0003 | *** | | |
| | Interaction | 5.89, 40.74 | 1.273 | 0.2916 | ns | | |
| **S9O** Beta | Dose | 1.48, 10.37 | 0.1338 | 0.8156 | ns | | |
| | Time | 5.53, 38.72 | 2.622 | 0.0346 | * | | |
| | Interaction | 5.29, 36.61 | 1.005 | 0.4314 | ns | | |
| **S9R** Low Gamma | Dose | 1.98, 13.85 | 6.097 | 0.0128 | * | Vehicle vs 30.0 mg/kg | 60, 90, 100, 120, 160, 210, 230, 270 |
| | Time | 5.78, 40.49 | 4.233 | 0.0024 | ** | | |
| | Interaction | 5.76, 39.87 | 1.675 | 0.1548 | ns | | |
| **S9U** High Gamma | Dose | 1.72, 12.01 | 2.399 | 0.1374 | ns | | |
| | Time | 5.44, 38.10 | 10.18 | <0.0001 | **** | | |
| | Interaction | 5.20, 35.97 | 1.499 | 0.213 | ns | | |
| **CP-101,606** | | | | | | | |
| **Ovx** | | | | | | | |
| **S10A** Delta | Dose | 1.76, 14.06 | 0.0807 | 0.9021 | ns | | |
| | Time | 4.88, 39.07 | 2.821 | 0.0296 | * | | |
| | Interaction | 5.75, 39.39 | 0.8943 | 0.5052 | ns | | |
| **S10C** Theta | Dose | 1.19, 9.541 | 0.5603 | 0.5021 | ns | | |
| | Time | 3.84, 30.72 | 2.537 | 0.0621 | ns | | |
| | Interaction | 5.12, 35.08 | 1.055 | 0.4024 | ns | | |
| **S10E** Alpha | Dose | 1.64, 13.10 | 0.1106 | 0.8587 | ns | | |
| | Time | 3.53, 28.25 | 3.992 | 0.0134 | * | | |
| | Interaction | 5.18, 35.50 | 0.8602 | 0.5203 | ns | | |
| **S10G** Sigma | Dose | 2.31, 18.44 | 1.448 | 0.2615 | ns | | |
| | Time | 4.47, 35.76 | 9.987 | <0.0001 | **** | | |
| | Interaction | 5.52, 37.84 | 1.075 | 0.3922 | ns | | |
| **S10I** Beta | Dose | 2.14, 17.08 | 0.1858 | 0.8451 | ns | | |
| | Time | 5.39, 43.13 | 3.871 | 0.0046 | * | | |
| | Interaction | 5.62, 38.52 | 1.146 | 0.3543 | ns | | |
| **S10K** Low Gamma | Dose | 2.14, 17.08 | 0.1858 | 0.8451 | ns | | |
| | Time | 5.39, 43.13 | 3.871 | 0.0046 | ** | | |
| | Interaction | 5.62, 38.52 | 1.146 | 0.3543 | ns | | |
| **S10M** High Gamma | Dose | 2.13, 17.00 | 1.302 | 0.2993 | ns | | |
| | Time | 4.81, 38.51 | 8.392 | <0.0001 | **** | | |
| | Interaction | 5.70, 39.11 | 0.8913 | 0.5067 | ns | | |
| **Ovx+E** | | | | | | | |
| **S10B** Delta | Dose | 2.48, 19.80 | 7.302 | 0.0027 | ** | Vehicle vs. 3.0 mg/kg<br>Vehicle vs. 30.0 mg/kg | 230<br>10, 20, 120 |
| | Time | 5.24, 41.94 | 6.611 | 0.0001 | *** | | |
| | Interaction | 6.62, 50.02 | 1.099 | 0.3775 | ns | | |

| | | | | | | | |
|---|---|---|---|---|---|---|---|
| **S10D** Theta | Dose | 2.68, 21.45 | 2.987 | 0.0586 | ns | | |
| | Time | 5.26, 42.05 | 3.016 | 0.019 | * | | |
| | Interaction | 6.54, 49.42 | 1.268 | 0.2874 | ns | | |
| **S10F** Alpha | Dose | 1.73, 13.82 | 2.463 | 0.1265 | ns | | |
| | Time | 4.54, 36.28 | 7.039 | 0.0002 | *** | | |
| | Interaction | 6.16, 46.57 | 1.123 | 0.3639 | ns | | |
| **S10H** Sigma | Dose | 2.00, 16.00 | 19.90 | <0.0001 | **** | Vehicle vs. 3.0 mg/kg<br>Vehicle vs 10.0 mg/kg<br>Vehicle vs. 30.0 mg/kg | -40, 270<br>20, 40, 50, 80, 170<br>30-50, 70-170, 250, 270 |
| | Time | 5.41, 43.29 | 9.541 | <0.0001 | **** | | |
| | Interaction | 6.18, 46.73 | 1.822 | 0.1133 | ns | | |
| **S10J** Beta | Dose | 2.06, 16.50 | 9.344 | 0.0018 | ** | Vehicle vs. 3.0 mg/kg<br>Vehicle vs. 30.0 mg/kg | -80, -20, 190, 270<br>50, 70, 80, 150, 160, 270 |
| | Time | 5.21, 41.65 | 3.637 | 0.0074 | ** | | |
| | Interaction | 6.27, 47.39 | 1.453 | 0.2128 | ns | | |
| **S10L** Low Gamma | Dose | 1.47, 11.74 | 0.7040 | 0.4719 | ns | | |
| | Time | 6.12, 48.96 | 4.991 | 0.0004 | *** | | |
| | Interaction | 6.44, 48.63 | 1.119 | 0.3659 | ns | | |
| **S10N** High Gamma | Dose | 1.32, 10.55 | 1.942 | 0.194 | ns | | |
| | Time | 4.47, 35.78 | 10.98 | <0.0001 | **** | | |
| | Interaction | 6.10, 46.07 | 1.413 | 0.2295 | ns | | |

Table S3. Statistics for Olanzapine and SBI-0646535 timecourse graphs.

| Timecourse Statistics | | | | | | | |
|---|---|---|---|---|---|---|---|
| **Figure** | **Source of Variation** | **DF** | **F** | **p** | ***** | ***Post hoc* results** | **Significant Timepoints (10-minute bin)** |
| **Olanzapine** | | | | | | | |
| **Ovary-Intact** | | | | | | | |
| **S11A** Delta | Dose | 2.58, 20.63 | 0.9685 | 0.4156 | ns | | |
| | Time | 4.56, 36.49 | 7.451 | <0.0001 | **** | | |
| | Interaction | 4.57, 33.92 | 0.9763 | 0.4412 | ns | | |
| **S11D** Theta | Dose | 3.34, 26.69 | 10.22 | <0.0001 | **** | Vehicle vs. 0.3 mg/kg | 30, 60, 110, 104, 150, 200 |
| | Time | 5.30, 42.39 | 13.74 | <0.0001 | **** | Vehicle vs. 0.56 mg/kg | 30-50, 110, 130, 170 |
| | Interaction | 5.92, 43.87 | 2.274 | 0.0543 | ns | Vehicle vs. 1.0 mg/kg | 30, 60, 80, 90, 140, 190 |
| | | | | | | Vehicle vs. 1.8 mg/kg | 20-40, 60-80, 110, 120, 140, 160, 230, 240 |
| **S11G** Alpha | Dose | 2.55, 20.36 | 0.8886 | 0.449 | ns | | |
| | Time | 4.15, 33.20 | 2.760 | 0.0421 | * | | |
| | Interaction | 5.50, 40.80 | 1.167 | 0.3423 | ns | | |
| **S11J** Sigma | Dose | 2.79, 22.31 | 1.725 | 0.193 | ns | | |
| | Time | 5.01, 40.05 | 10.97 | <0.0001 | **** | | |
| | Interaction | 5.50, 40.78 | 1.104 | 0.3752 | ns | | |
| **S11M** Beta | Dose | 2.13, 17.04 | 0.8858 | 0.4366 | ns | | |
| | Time | 3.80, 30.39 | 6.360 | 0.0009 | *** | | |
| | Interaction | 4.26, 31.57 | 1.046 | 0.4018 | ns | | |
| **S11P** Low Gamma | Dose | 2.63, 21.03 | 6.943 | 0.0027 | ** | Vehicle vs. 0.3 mg/kg | 40, 50, 90, 140, 170 |
| | Time | 5.10, 40.77 | 14.91 | <0.0001 | **** | Vehicle vs. 0.56 mg/kg | 90, 110, 120, 250 |
| | Interaction | 6.15, 45.64 | 1.606 | 0.1657 | ns | Vehicle vs. 1.0 mg/kg | 40, 90-120 |
| | | | | | | Vehicle vs. 1.8 mg/kg | 10-30, 50, 60, 110, 120, 140, 170 |
| **S11S** High Gamma | Dose | 2.76, 22.06 | 7.408 | 0.0016 | ** | Vehicle vs. 0.3 mg/kg | 40, 50, 90, 140, 170, 190 |
| | Time | 5.71, 45.67 | 25.09 | <0.0001 | **** | Vehicle vs. 0.56 mg/kg | 90, 110-130 |
| | Interaction | 6.57, 48.75 | 1.447 | 0.2119 | ns | Vehicle vs. 1.0 mg/kg | 0, 40, 60, 90-120, 150 |
| | | | | | | Vehicle vs. 1.8 mg/kg | 10-30, 50, 60, 110, 130, 140, 160 |
| **Ovx** | | | | | | | |
| **S11B** Delta | Dose | 2.53, 20.25 | 2.673 | 0.0826 | ns | | |
| | Time | 4.80, 38.42 | 4.606 | 0.0024 | ** | | |
| | Interaction | 4.99, 35.43 | 1.603 | 0.1849 | ns | | |
| **S11E** Theta | Dose | 2.51, 20.10 | 6.377 | 0.0047 | ** | Vehicle vs. 0.3 mg/kg | 20, 50-70, 120 |
| | Time | 3.79, 30.35 | 9.477 | <0.0001 | **** | Vehicle vs. 0.56 mg/kg | 50+60, 100, 170 |
| | Interaction | 4.48, 31.81 | 1.776 | 0.1523 | ns | Vehicle vs. 1.0 mg/kg | 10, 20, 40, 60, 90, 150, 170 |
| | | | | | | Vehicle vs. 1.8 mg/kg | 0-60, 100 |
| **S11H** Alpha | Dose | 2.52, 20.18 | 1.260 | 0.1402 | ns | | |
| | Time | 3.91, 31.28 | 1.878 | 0.3111 | ns | | |
| | Interaction | 4.99, 35.43 | 1.193 | 0.3325 | ns | | |
| **S11K** Sigma | Dose | 2.85, 22.80 | 0.5619 | 0.6371 | ns | | |
| | Time | 4.21, 33.71 | 9.612 | <0.0001 | **** | | |
| | Interaction | 4.35, 30.91 | 1.117 | 0.3688 | ns | | |
| **S11N** Beta | Dose | 2.51, 20.06 | 1.724 | 0.1992 | ns | | |
| | Time | 3.71, 29.69 | 2.980 | 0.038 | * | | |
| | Interaction | 3.53, 25.09 | 1.219 | 0.3263 | ns | | |
| **S11Q** Low Gamma | Dose | 2.16, 17.26 | 2.038 | 0.1581 | ns | | |
| | Time | 4.18, 33.46 | 8.535 | <0.0001 | **** | | |
| | Interaction | 4.17, 29.60 | 1.552 | 0.2116 | ns | | |
| **S11T** High Gamma | Dose | 2.09, 16.75 | 1.479 | 0.2565 | ns | | |
| | Time | 4.26, 34.04 | 18.28 | <0.0001 | **** | | |
| | Interaction | 4.29, 30.46 | 1.468 | 0.2341 | ns | | |
| **Ovx+E** | | | | | | | |
| **S11C** Delta | Dose | 4.00, 32.00 | 1.816 | 0.1501 | ns | Vehicle vs. 1.0 mg/kg | 240 |
| | Time | 42.0, 336.0 | 8.574 | <0.0001 | **** | | |

| | | | | | | | |
|---|---|---|---|---|---|---|---|
| | Interaction | 168, 1195 | 1.564 | <0.0001 | **** | | |
| **S11F** Theta | Dose | 4.00, 32.00 | 12.77 | <0.0001 | **** | Vehicle vs. 0.3 mg/kg<br>Vehicle vs. 0.56 mg/kg<br>Vehicle vs. 1.0 mg/kg<br>Vehicle vs. 1.8 mg/kg | 40, 80<br>10<br>-80, 10, 30, 210<br>10-40, 60, 90, 160, 210, 230, 260-300 |
| | Time | 42.0, 336.0 | 6.056 | <0.0001 | **** | | |
| | Interaction | 168, 1195 | 1.711 | <0.0001 | **** | | |
| **S11I** Alpha | Dose | 4.00, 32.00 | 3.822 | 0.0119 | * | Vehicle vs. 0.3 mg/kg<br>Vehicle vs. 1.0 mg/kg<br>Vehicle vs. 1.8 mg/kg | -70, -30, 40<br>100, 130<br>-10, 210, 220, 260 |
| | Time | 42.0, 336.0 | 4.689 | <0.0001 | **** | | |
| | Interaction | 168, 1195 | 1.313 | 0.0073 | ** | | |
| **S11L** Sigma | Dose | 4.00, 32.00 | 1.504 | 0.2245 | ns | | |
| | Time | 42.0, 336.0 | 8.248 | <0.0001 | **** | | |
| | Interaction | 168, 1195 | 1.132 | 0.1344 | ns | | |
| **S11O** Beta | Dose | 4.00, 32.00 | 2.861 | 0.0392 | * | Vehicle vs. 0.3 mg/kg<br>Vehicle vs. 1.8 mg/kg | 300<br>80 |
| | Time | 42.0, 336.0 | 6.855 | <0.0001 | **** | | |
| | Interaction | 168, 1195 | 1.484 | 0.0002 | *** | | |
| **S11R** Low Gamma | Dose | 4.00, 32.00 | 5.214 | 0.0024 | ** | Vehicle vs. 0.3 mg/kg<br>Vehicle vs. 1.0 mg/kg<br>Vehicle vs. 1.8 mg/kg | 60<br>190, 240<br>110, 170 |
| | Time | 42.0, 336.0 | 9.443 | <0.0001 | **** | | |
| | Interaction | 168, 1195 | 1.184 | 0.066 | ns | | |
| **S11U** High Gamma | Dose | 4, 32 | 4.78 | 0.0039 | ** | Vehicle vs. 1.0 mg/kg<br>Vehicle vs. 1.8 mg/kg | 190, 240<br>20, 90 |
| | Time | 42, 336 | 16.14 | <0.0001 | **** | | |
| | Interaction | 168, 1195 | 1.126 | 0.144 | Ns | | |
| **SBI-0646535** | | | | | | | |
| **Ovary-Intact** | | | | | | | |
| **S12A** Delta | Dose | 1.97, 13.77 | 1.500 | 0.2573 | ns | | |
| | Time | 3.47, 24.26 | 1.520 | 0.2315 | ns | | |
| | Interaction | 4.34, 30.06 | 1.295 | 0.2939 | ns | | |
| **S12D** Theta | Dose | 2.18, 15.25 | 0.6970 | 0.525 | ns | | |
| | Time | 5.10, 35.68 | 1.569 | 0.1929 | ns | | |
| | Interaction | 6.25, 43.22 | 1.009 | 0.4336 | ns | | |
| **S12G** Alpha | Dose | 2.70, 18.91 | 1.397 | 0.2743 | ns | | |
| | Time | 5.18, 36.28 | 3.159 | 0.0171 | * | | |
| | Interaction | 6.03, 41.74 | 1.256 | 0.2983 | ns | | |
| **S12J** Sigma | Dose | 1.86, 13.05 | 1.091 | 0.3603 | ns | | |
| | Time | 5.62, 39.34 | 5.504 | 0.0004 | *** | | |
| | Interaction | 6.21, 42.95 | 1.043 | 0.4123 | ns | | |
| **S12M** Beta | Dose | 2.42, 16.96 | 2.643 | 0.0921 | ns | | |
| | Time | 4.20, 29.43 | 3.513 | 0.0171 | * | | |
| | Interaction | 5.10, 35.31 | 1.030 | 0.4162 | ns | | |
| **S12P** Low Gamma | Dose | 2.37, 16.61 | 1.421 | 0.2713 | ns | | |
| | Time | 5.09, 35.62 | 5.113 | 0.0012 | ** | | |
| | Interaction | 6.09, 42.15 | 1.433 | 0.2241 | ns | | |
| **S12S** High Gamma | Dose | 2.11, 14.74 | 0.2068 | 0.826 | ns | | |
| | Time | 5.07, 35.51 | 8.109 | <0.0001 | **** | | |
| | Interaction | 6.06, 41.91 | 1.632 | 0.1618 | ns | | |
| **Ovx** | | | | | | | |
| **S12B** Delta | Dose | 2.21, 17.64 | 1.728 | 0.2048 | ns | | |
| | Time | 4.46, 35.70 | 1.314 | 0.282 | ns | | |
| | Interaction | 6.26, 44.59 | 1.339 | 0.2588 | ns | | |
| **S12E** Theta | Dose | 1.37, 10.93 | 2.108 | 0.1739 | ns | | |
| | Time | 2.49, 19.93 | 1.553 | 0.2347 | ns | | |
| | Interaction | 4.17, 29.72 | 1.129 | 0.3626 | ns | | |
| **S12H** Alpha | Dose | 1.22, 9.738 | 1.063 | 0.3441 | ns | | |
| | Time | 2.70, 21.56 | 1.330 | 0.2898 | ns | | |
| | Interaction | 3.97, 28.25 | 0.8152 | 0.5252 | ns | | |

| | | | | | |
|---|---|---|---|---|---|
| **S12K** Sigma | Dose | 1.68, 13.45 | 0.5788 | 0.5455 | ns |
| | Time | 3.57, 28.56 | 5.838 | 0.002 | ** |
| | Interaction | 3.98, 28.38 | 0.8951 | 0.4794 | ns |
| **S12N** Beta | Dose | 1.32, 10.58 | 0.5041 | 0.5437 | ns |
| | Time | 3.74, 29.95 | 2.823 | 0.0453 | * |
| | Interaction | 4.11, 29.30 | 1.024 | 0.4125 | ns |
| **S12Q** Low Gamma | Dose | 2.70, 21.58 | 0.7719 | 0.5099 | ns |
| | Time | 5.58, 44.66 | 5.957 | 0.0002 | *** |
| | Interaction | 6.11, 43.56 | 0.9626 | 0.4626 | ns |
| **S12T** High Gamma | Dose | 1.86, 14.90 | 0.1395 | 0.8572 | ns |
| | Time | 5.30, 42.36 | 10.21 | <0.0001 | **** |
| | Interaction | 6.06, 43.20 | 1.213 | 0.3179 | ns |
| **Ovx+E** | | | | | |
| **S12C** Delta | Dose | 1.51, 12.11 | 2.521 | 0.1299 | ns |
| | Time | 4.64, 37.14 | 4.633 | 0.0027 | ** |
| | Interaction | 5.22, 36.62 | 1.268 | 0.2979 | ns |
| **S12F** Theta | Dose | 2.34, 18.75 | 0.1775 | 0.8692 | ns |
| | Time | 4.35, 34.79 | 0.8048 | 0.5393 | ns |
| | Interaction | 5.77, 40.47 | 0.8788 | 0.5158 | ns |
| **S12I** Alpha | Dose | 1.63, 13.00 | 0.5264 | 0.5665 | ns |
| | Time | 5.60, 44.82 | 2.813 | 0.023 | * |
| | Interaction | 6.15, 43.13 | 1.607 | 0.1671 | ns |
| **S12L** Sigma | Dose | 2.40, 19.20 | 1.087 | 0.3671 | ns |
| | Time | 4.63, 37.07 | 5.406 | 0.001 | ** |
| | Interaction | 5.79, 40.64 | 1.316 | 0.2729 | ns |
| **S12O** Beta | Dose | 2.05, 16.38 | 2.575 | 0.1056 | ns |
| | Time | 3.57, 28.55 | 1.475 | 0.2386 | ns |
| | Interaction | 4.85, 34.03 | 1.192 | 0.3338 | ns |
| **S12R** Low Gamma | Dose | 2.10, 16.77 | 3.065 | 0.0714 | ns |
| | Time | 4.47, 35.73 | 6.214 | 0.0005 | *** |
| | Interaction | 5.58, 39.13 | 1.291 | 0.2855 | ns |
| **S12U** High Gamma | Dose | 2.50, 20.02 | 0.8284 | 0.475 | ns |
| | Time | 4.97, 39.77 | 10.14 | <0.00011 | **** |
| | Interaction | 5.63, 39.49 | 1.162 | 0.3457 | ns |

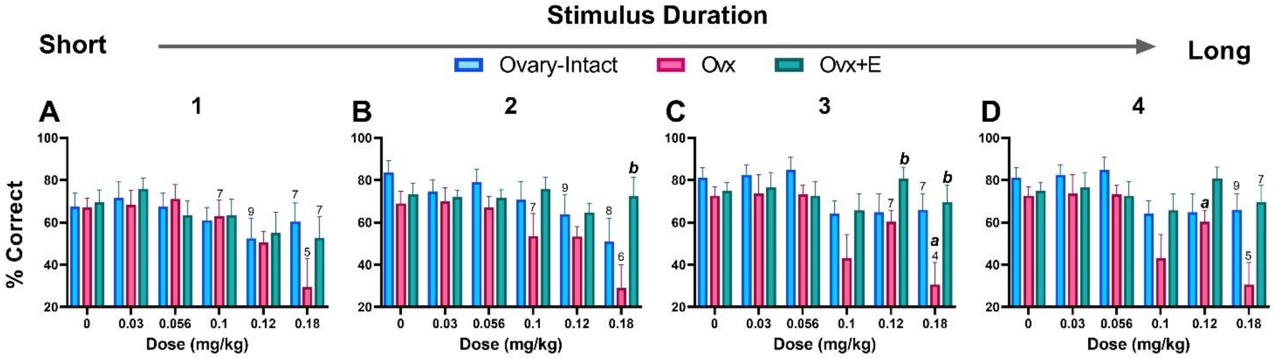

**Figure S1.** Ovx rats had lower accuracy compared to Ovx+E rats in the middle two stimulus durations. Bars depict group data as the mean (±SEM) % correct across each tested dose. At stimulus duration 1 (the shortest duration), there was a main effect of dose ($F_{3.59, 76.19} = 7.036$, p=0.001) but no effect of group or interaction (ps>0.05) (**A**). At stimulus duration 2, there was a main effect of dose $F_{3.10, 66.98} = 5.601$, p=0.0015) and group ($F_{2, 23} = 4.707$, p=0.0193) but no interaction (**B**). Ovx rats had lower accuracy compared to Ovx+E rats and experienced significant reductions relative to their respective vehicle condition at the 0.18 mg/kg dose. At stimulus duration 3, there was a main effect of dose ($F_{3.97, 84.06} = 6.823$; p<0.0001) and group ($F_{2, 23} = 5.650$; p=0.0101) but no interaction (**C**). Ovx rats had lower accuracy compared to Ovx+E rats at the 0.12 and 0.18 mg/kg dose, and decreased accuracy relative to their respective vehicle condition at the 0.18 mg/kg dose. Lastly, at stimulus duration 4 (the longest stimulus duration), there was a main effect of dose ($F_{3.44, 75.03} = 4.561$; p=0.0037) but no effect of group or interaction (ps>0.05) (**D**). Ovx rats displayed significant decreases relative to vehicle at the 0.12 mg/kg dose. Some rats were excluded due to omission of all trials within a stimulus duration; in those cases, the included n is depicted on the bar graph. *p*<0.05; *a* compared to the group's respective vehicle condition; *b* compared to the Ovx group. n=10 ovary-intact, 8 Ovx, and 8 Ovx+E rats.

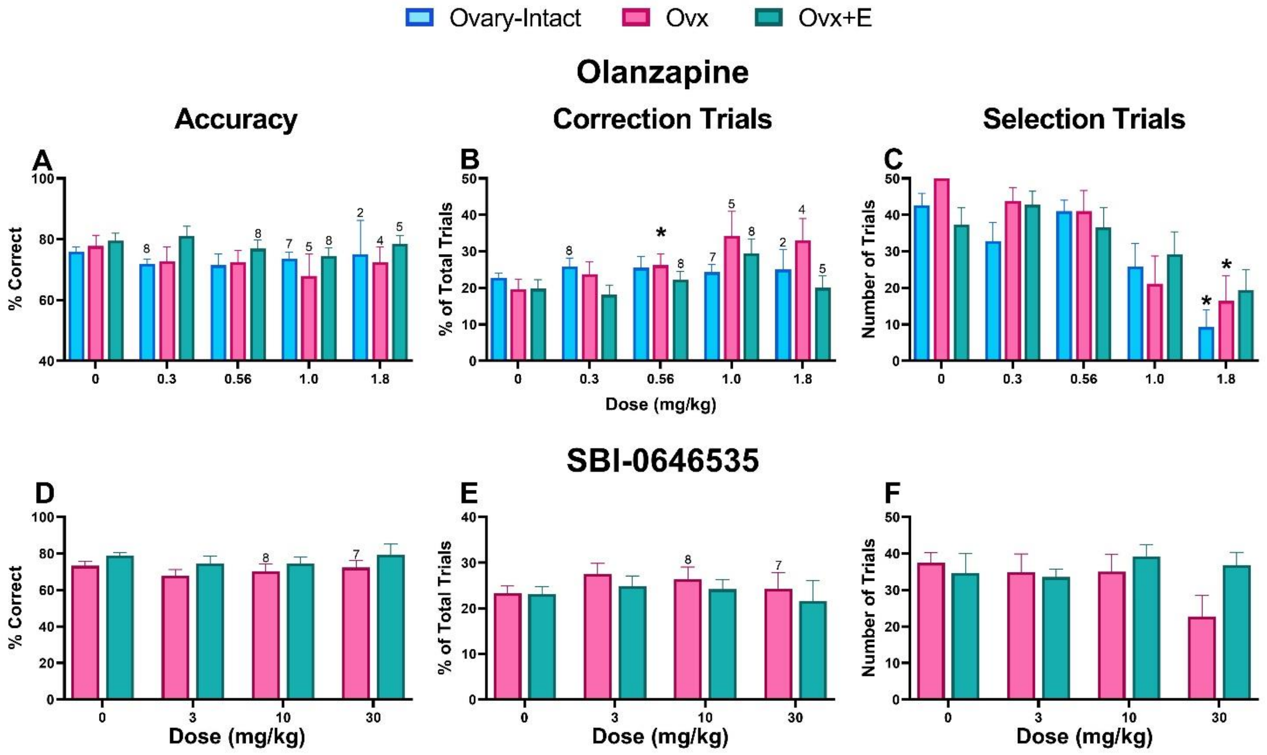

**Figure S2.** OLZ and SBI-0646535 did not affect accuracy on the PAL task in either group. There were no significant effects of OLZ on % correct (**A**). There was a main effect of dose ($F_{3.30, 56.83}$ = 4.436; p=0.0057) but no effect of group or interaction (ps>0.05) on % CTs, where only Ovx rats experienced a significant increase at the 0.56 mg/kg dose (**B**). Lastly, in line with the sedative profile of OLZ, there was a main effect of dose ($F_{3.16, 75.00}$ = 25.06; p<0.0001), but no effect of group or interaction (ps>0.05), on number of selection trials. The 1.8 mg/kg dose significantly reduced selection trials in ovary-intact and Ovx rats (**C**). SBI-0646535 did not significantly affect accuracy (**D**), correction trials (**E**), or selection trials (**F**). *p<0.05 compared to each group's respective vehicle condition. For OLZ, n= 11 ovary-intact, 8 Ovx, and 9 Ovx+E rats. For SBI-0646535, n=9 Ovx and 7 Ovx+E. However, not all rats performed at inclusion criteria (≥10 trials) on any given dose day; the included n is depicted on the bar graph if there were rats excluded.

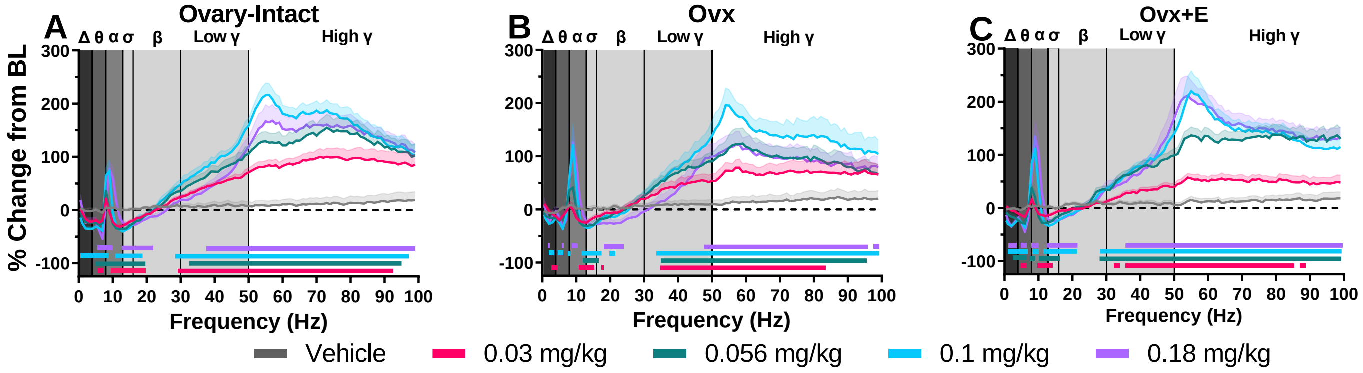

**Figure S3.** All tested MK-801 doses influence spectral frequencies across waveforms in each group. Data are expressed as a group mean (± SEM) presented in 1 Hz bins as the average percent change from baseline during the 30-90 minute post-dosing period. Gray vertical bars represent frequency bands (delta, Δ 0.5–4 Hz; theta, θ 4–8 Hz; alpha, α 8–13 Hz; sigma, σ 13–15 Hz; beta, β 13–30 Hz; low gamma, γ 30–50 Hz; high gamma, γ 50–100 Hz). All tested doses were examined within each individual group: ovary-intact (n=9-10) (**A**), Ovx (n=8-9) (**B**), and Ovx +E (n=12) rats (**C**). p<0.05; horizontal colored lines matching the respective dose color represent frequencies at which MK-801-treated groups were significantly different from vehicle-treated groups (**A–C**)

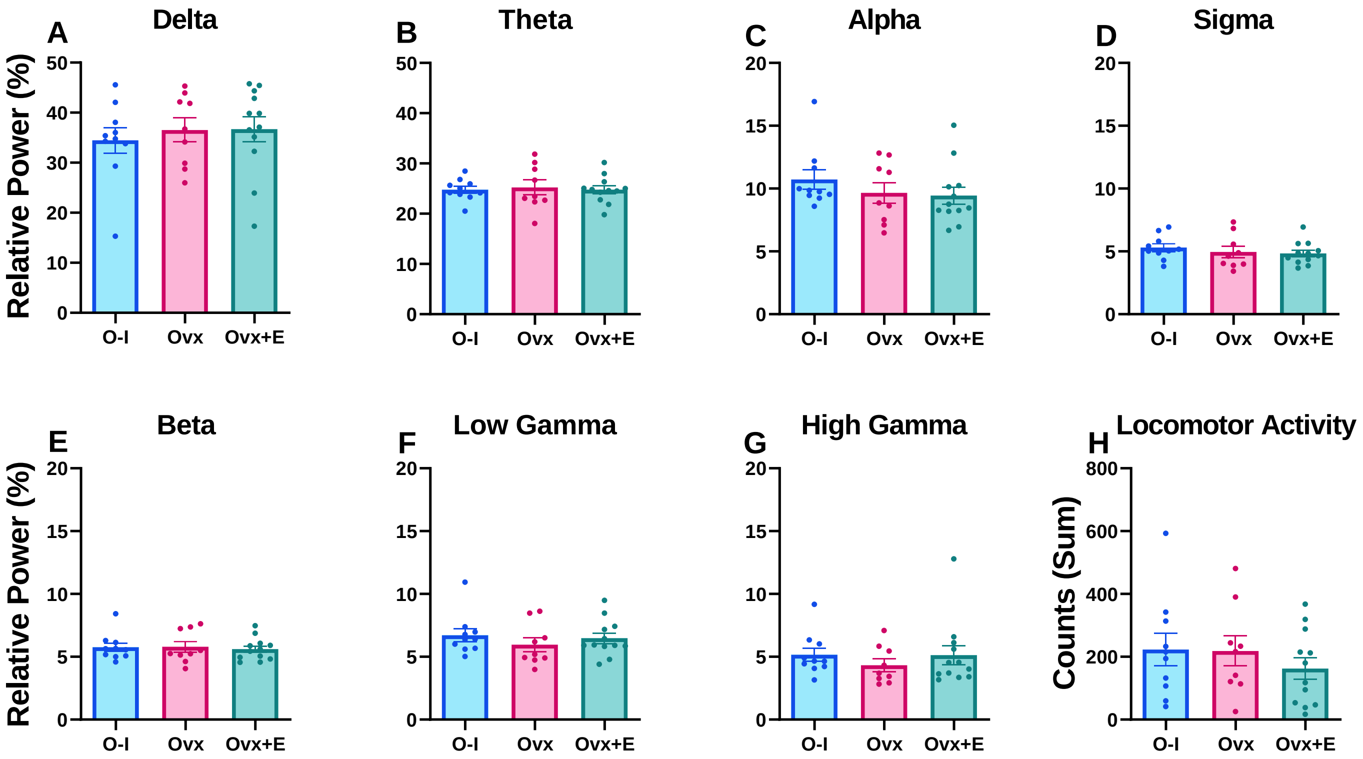

**Figure S4.** There were no group differences in relative power during the baseline period. qEEG data are shown as a group mean (±SEM) of relative power during the 90 minute baseline period on each individuals' respective vehicle day for MK-801. Locomotor activity is expressed as summed activity counts during the baseline period. Data are averaged across the delta (**A**), theta (**B**), alpha (**C**), sigma (**D**), beta (**E**), low gamma (**F**), high gamma (**G**) frequency bands and activity was summed (**H**) to compare O-I (n=9), Ovx (n=9) and Ovx+E (n=12) rats; circles represent individual datapoints.

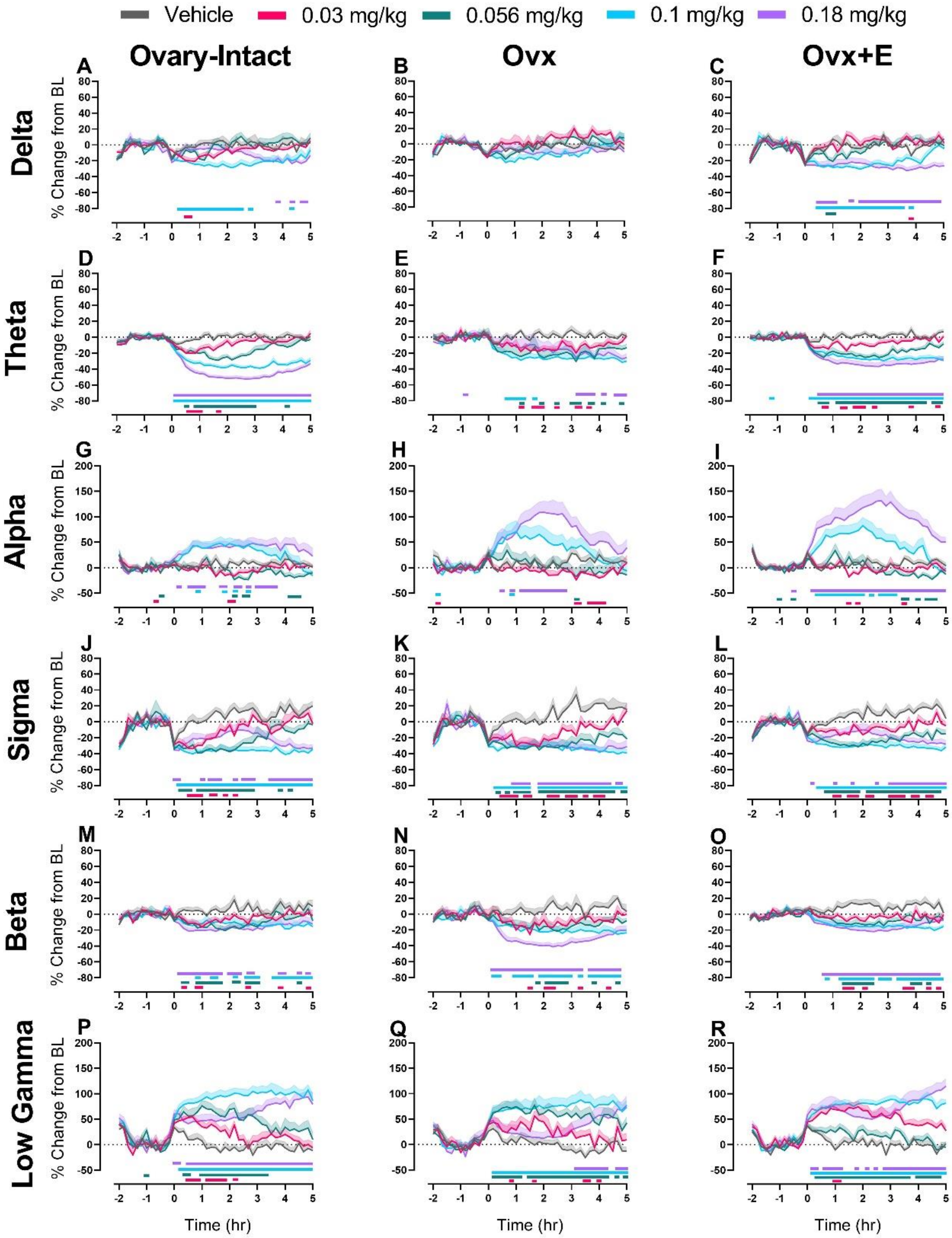

**Figure S5. MK-801 significantly affects all spectral power bands in each tested group.** Time course effects of MK-801 are displayed as group means (±SEM) as the percent change from the 90-minute baseline in 10-minute bins across the 7-hour recording period on delta (**A-C**), theta (**D-F**), alpha (**G-I**), sigma (**J-L**), beta (**M-O**), and low gamma (**P-R**) in O-I, Ovx, and Ovx+E rats. MK-801 was administered at time point 0. On the x-axis, -2 corresponds to ZT 0 and 5 corresponds to ZT 7. Horizontal colored lines matching the respective dose color represent the 10-minute bins at which MK-801 treatment was significantly different from vehicle treatment. $p<0.05$, n= 9-10 O-I, 8-9 Ovx, and 12 Ovx+E rats.

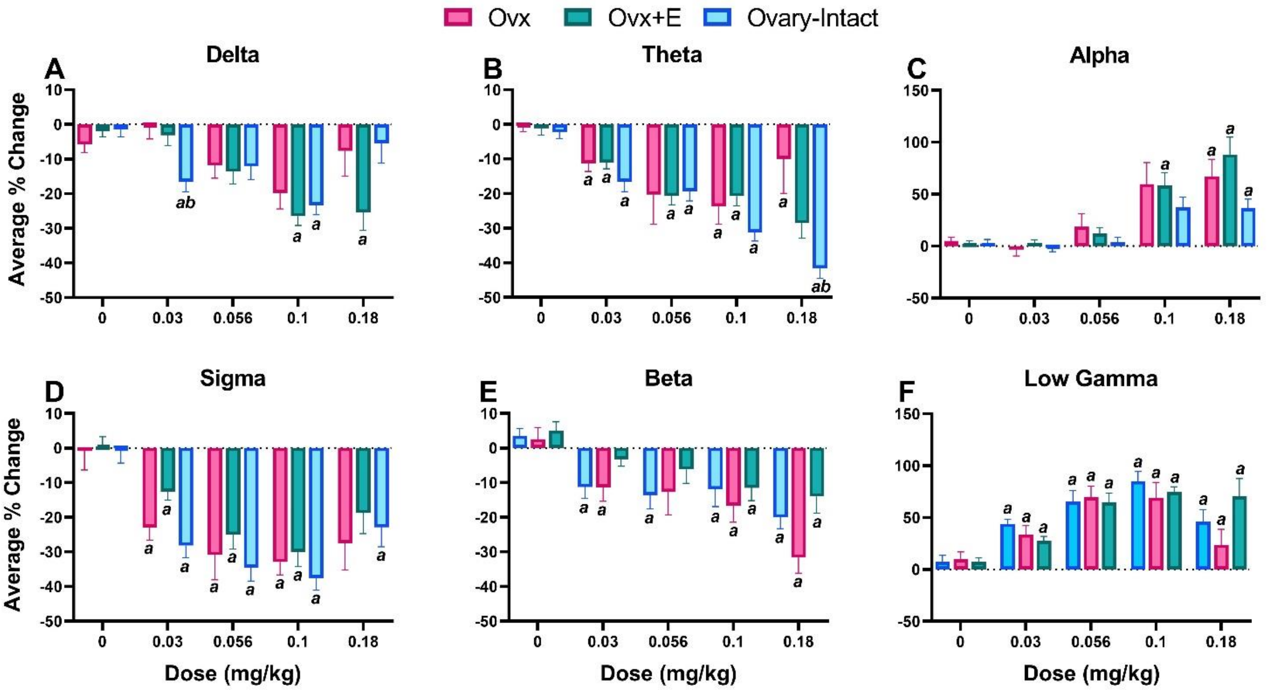

**Figure S6.** Significant MK-801-induced changes were observed in all waveforms with significant group differences in the delta and theta waveforms. For direct group comparisons in response to MK-801, each individual's percent change from baseline in the 30-90 min post-dosing period was averaged and graphed as a group mean ±SEM to assess the effects of MK-801 administration on delta (**A**), theta (**B**), alpha (**C**), sigma (**D**), beta (**E**), and low gamma power (**F**). A two-way ANOVA was run for each frequency band. For delta, there was a main effect of dose ($F_{2.65, 71.41} = 13.06$, $p<0.0001$) and a dose x group interaction ($F_{8, 108} = 3.362$, $p=0.0018$). O-I (0.03 and 0.1 mg/kg) and Ovx+E (0.1 and 0.18 mg/kg) rats exhibited significant decreases, and O-I rats showed a significantly greater decrease than Ovx rats at the 0.03 mg/kg dose (**A**). For theta, there was a main effect of dose ($F_{2.63, 71.02} = 24.53$, $p<0.0001$) and a dose x group interaction ($F_{8, 108} = 3.260$ $p=0.0023$). All groups showed significant decreases several doses indicated above, and O-I rats had greater decreases than Ovx rats at the 0.18 mg/kg dose (**B**). For alpha, there was a main effect of dose ($F_{2.65, 71.55} = 27.72$, $p<0.0001$), where all groups showed significant, dose-dependent increases at the 0.18 mg/kg dose (**C**). For sigma, there was a main effect of dose ($F_{3.16, 85.31} = 32.19$, $p<0.0001$), with significant decreases at multiple doses shown in all groups (**D**). For beta, there was a man effect of dose ($F_{3.34, 90.21} = 21.53$, $p<0.0001$), with significant decreases at multiple doses shown in all groups (**E**). Lastly, for low gamma, there was a main effect of dose ($F_{2.22, 59.81} = 24.26$, $p<0.0001$) where all groups showed increases in low gamma power at all tested doses (**F**). $p<0.05$, *a* compared to the group's respective vehicle condition; *b* compared to the Ovx group.

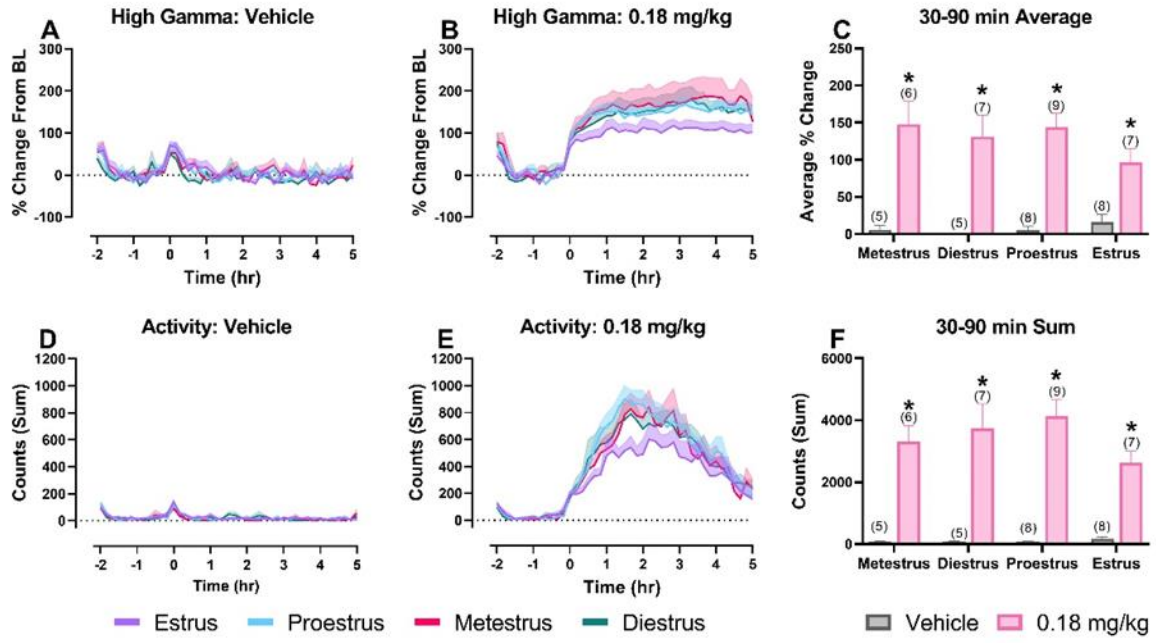

**Figure S7.** There was no significant effect of estrous cycle phase following administration of vehicle or 0.18 mg/kg MK-801. Time course effects of the different phases following vehicle (**A**) and MK-801 (**B**) on high gamma power are displayed as group means ±SEM of the percent change from 90-minute baseline in 10-minute bins across the 7-hour recording period. Time course effects of vehicle (**D**) and MK-801 (**E**) on locomotor activity are displayed as group means ±SEM of the summed activity counts in 10-minute bins across the 7-hour recording period. Averages of the high gamma power % change from baseline (**C**) and summed locomotor activity (**F**) were evaluated for vehicle and 0.18 mg/kg in the 30-90 minute post-dosing period. $p<0.05$; * significantly different from the vehicle condition in the respective estrous cycle phase.

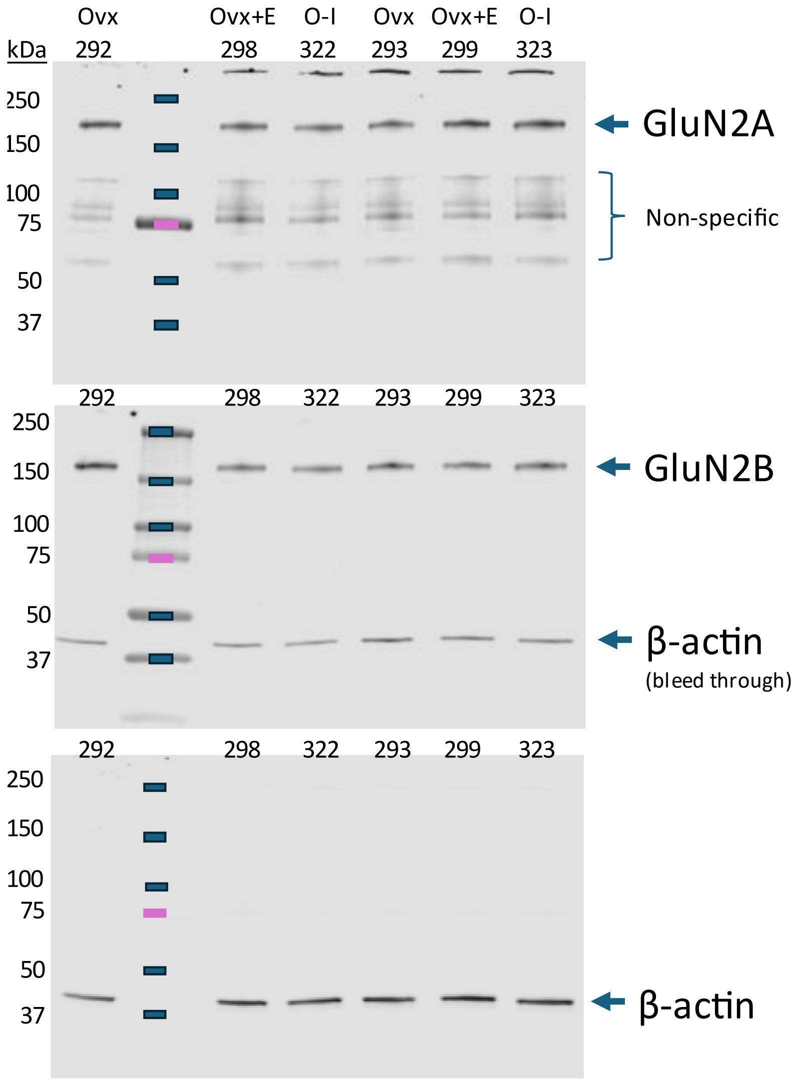

**Figure S8:** Additional representative Western blots of estrogen regulated expression of GluN2A and 2B. Cortical synaptoneurosomes were solubilized in SDS-sample buffer and subjected to SDS-PAGE and Western blot analysis using antibodies against GluN2A, GluN2B, and the house keeping protein beta-actin. Treatment is indicated above. Numbers under treatment are mouse ID. Molecular weight standards (Biorad) were run on the gel in the second to left land. Molecular weight is noted in kilodaltons on the left of each Western blot. Note, GluN2A, GluN2B, and Beta-actin run approximately at their predicted molecular weight (arrow) of 190 kD, 170kD, and 43 kDa, respectively.

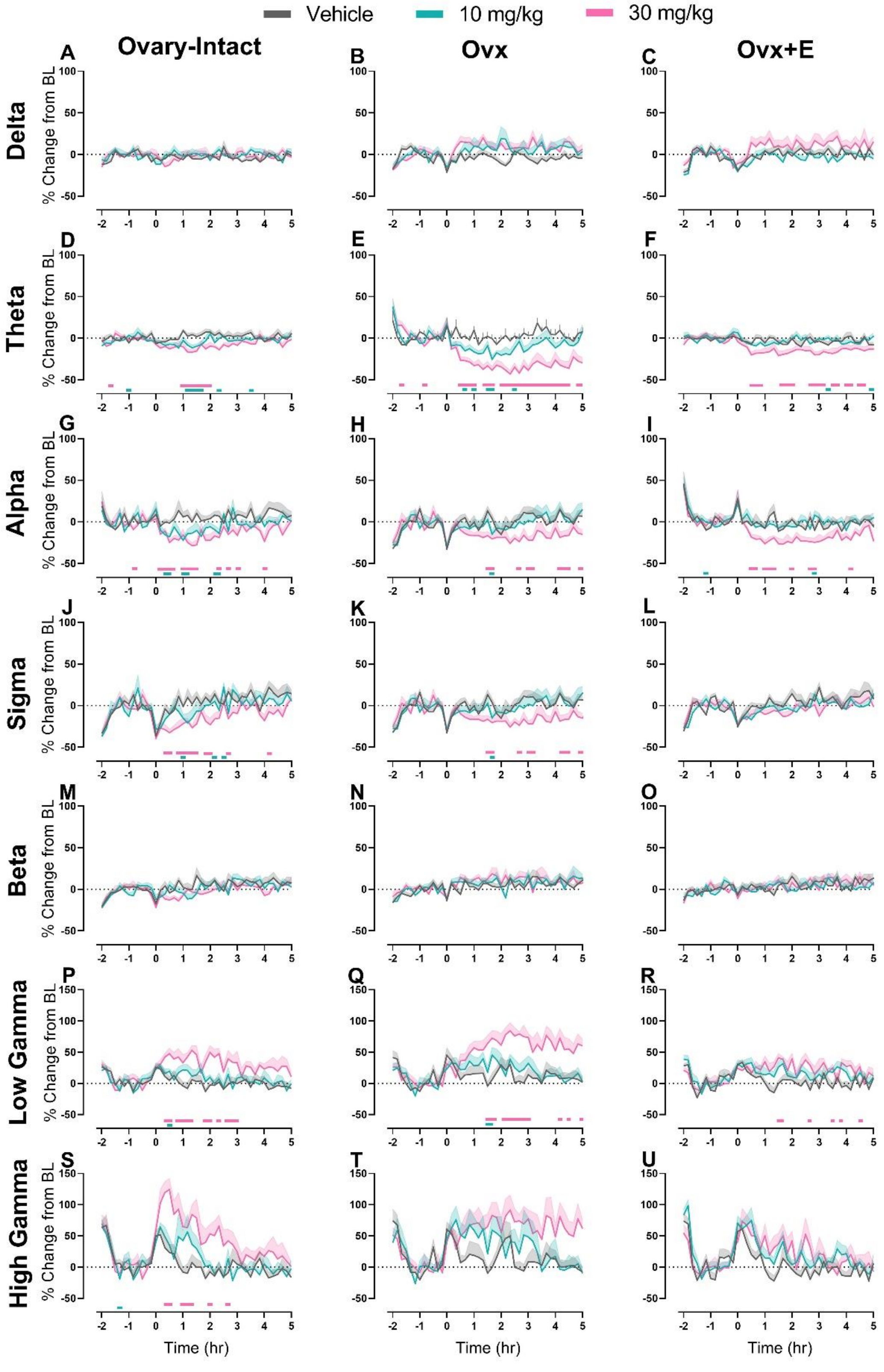

**Figure S9.** PEAQX significantly affected multiple spectral power bands in each tested group. Time course effects of PEAQX are displayed as group means ±SEM as the percent change from the 90-minute baseline in 10-minute bins across the 7-hour recording period on delta (**A-C**), theta (**D-F**), alpha (**G-I**), sigma (**J-L**), beta (**M-O**), low gamma (**P-R**), and high gamma (**S-U**) in ovary-intact, Ovx, and Ovx+E rats. PEAQX was administered at time point 0. On the x-axis, -2 corresponds to ZT 0 and 5 corresponds to ZT 7. Horizontal colored lines matching the respective dose color represent the 10-minute bins at which PEAQX treatment was significantly different from vehicle treatment. $p<0.05$, n= 7-8 ovary-intact, 8 Ovx, and 8 Ovx+E rats.

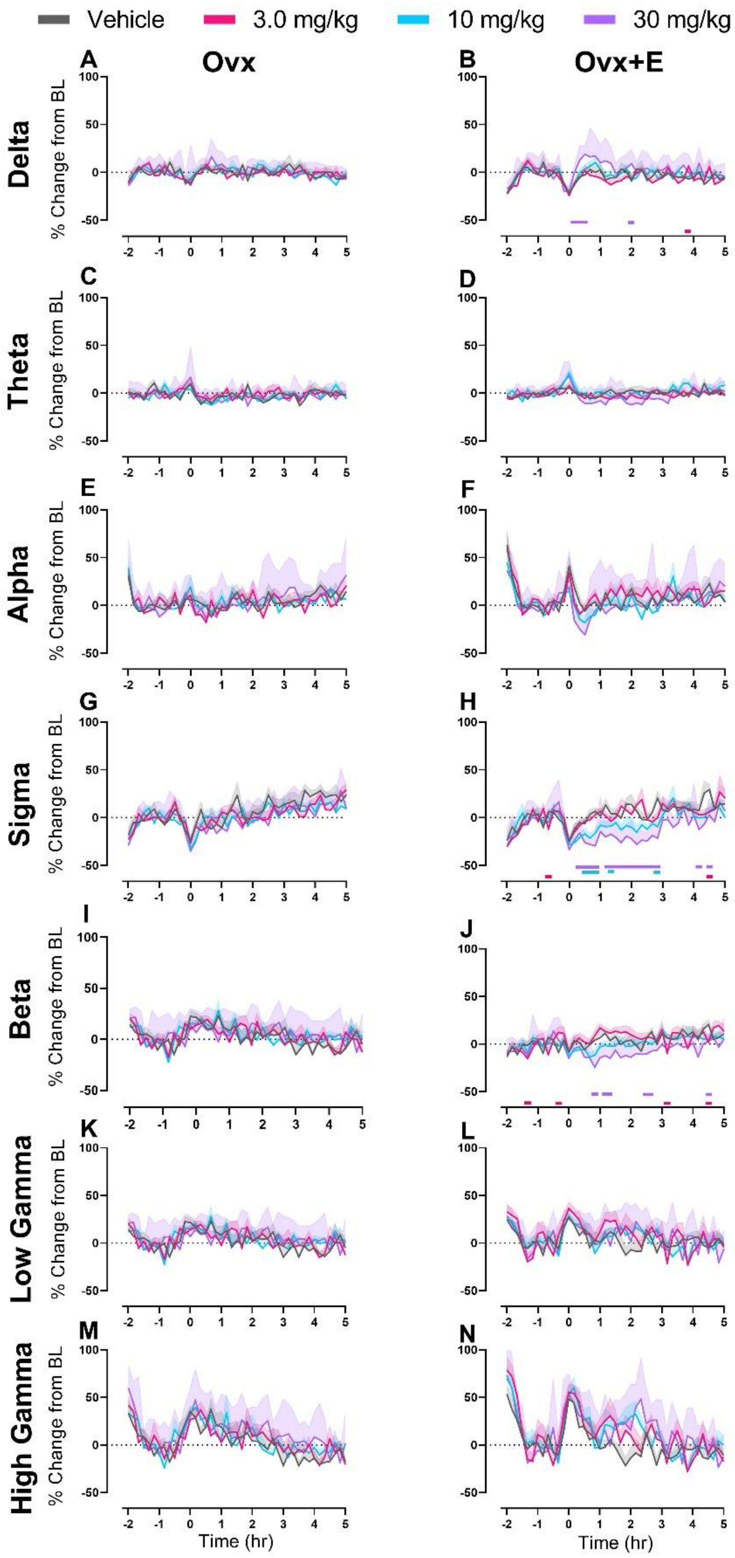

**Figure S10.** CP-101,606 only affected sigma and beta power in Ovx+E rats. Time course effects of CP-101,606 are displayed as group means ±SEM as the percent change from the 90-minute baseline in 10-minute bins across the 7-hour recording period on delta (**A,B**), theta (**C,D**), alpha (**E,F**), sigma (**G,H**), beta (**I,J**), low gamma (**K,L**), and high gamma (**M,N**) in ovary-intact, Ovx, and Ovx+E rats. CP-101,606 was administered at time point 0. On the x-axis, -2 corresponds to ZT 0 and 5 corresponds to ZT 7. Horizontal colored lines matching the respective dose color represent the 10-minute bins at which CP-101,606 treatment was significantly different from vehicle treatment. $p<0.05$, n= 8-9 Ovx and 8-9 Ovx+E rats.

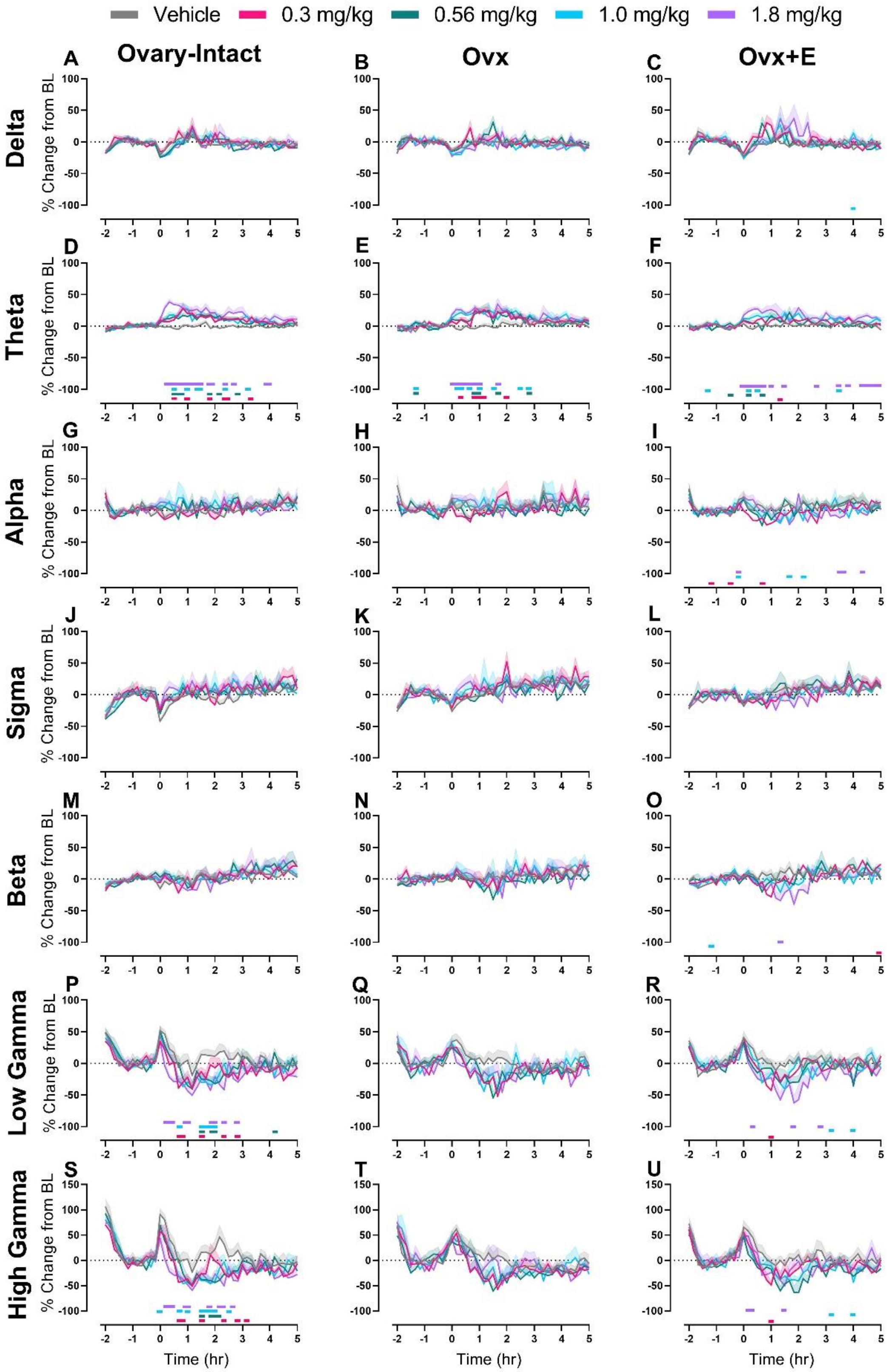

**Figure S11.** OLZ significantly affected distinct waveforms over time within each group. Time course effects of olanzapine are displayed as group means (±SEM) as the percent change from the 90-minute baseline in 10-minute bins across the 7-hour recording period on delta (**A-C**), theta (**D-F**), alpha (**G-I**), sigma (**J-L**), beta (**M-O**), low gamma (**P-R**), and high gamma (**S-U**) in O-I, Ovx, and Ovx+E rats. OLZ was administered at time point 0. On the x-axis, -2 corresponds to ZT 0 and 5 corresponds to ZT 7. Horizontal colored lines matching the respective dose color represent the 10-minute bins at which OLZ treatment was significantly different from vehicle treatment. $p<0.05$, $n= 9$ per group.

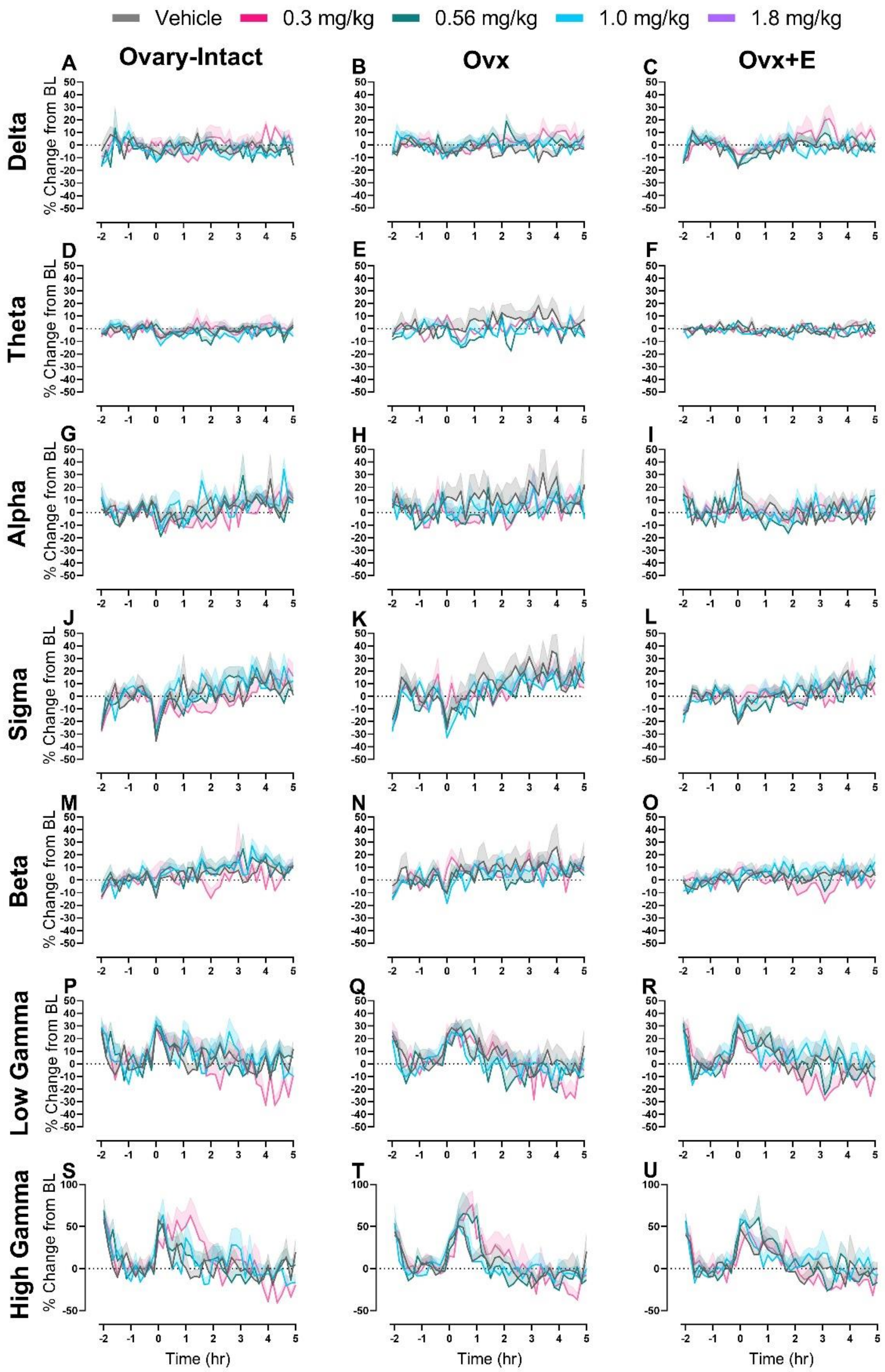

**Figure S12.** SBI-0646535 produced no significant effects on any waveforms in each tested group. Time course effects of SBI-0646535 are displayed as group means (±SEM) as the percent change from the 90-minute baseline in 10-minute bins across the 7-hour recording period on delta (**A-C**), theta (**D-F**), alpha (**G-I**), sigma (**J-L**), beta (**M-O**), low gamma (**P-R**), and high gamma (**S-U**) in O-I, Ovx, and Ovx+E rats. SBI-0646535 was administered at time point 0. On the x-axis, -2 corresponds to ZT 0 and 5 corresponds to ZT 7. Horizontal colored lines matching the respective dose color represent the 10-minute bins at which SBI-0646535 treatment was significantly different from vehicle treatment. $p<0.05$, n= 8 ovary-intact, 7-9 Ovx, and 7-9 Ovx+E rats.

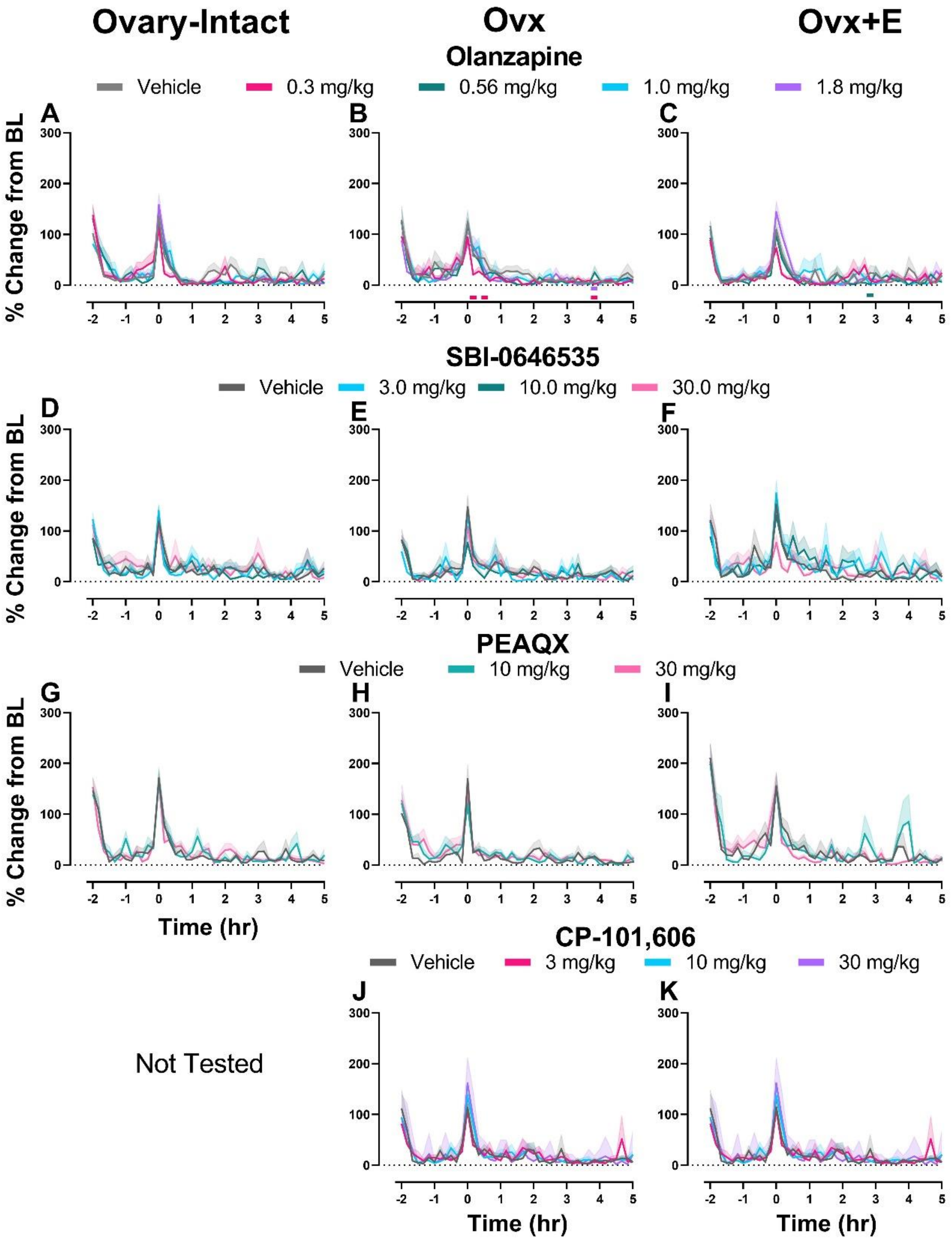

**Figure S13.** Only OLZ significantly affected locomotor activity when administered alone. Time course effects are displayed as group means (±SEM) as the summed activity counts in 10-minute bins across the 7-hour recording period following administration of OLZ (**A-C**), SBI-0646535 (**D-F**), PEAQX (**G-I**), and CP-101,606 sigma (**J,K**) to O-I, Ovx, and Ovx+E rats. Compounds were administered at time point 0. On the x-axis, -2 corresponds to ZT 0 and 5 corresponds to ZT 7. Significant effects of dose on locomotor activity were only found in Ovx (main effect of dose, $F_{3.01, 24.08} = 4.922$, p=0.0083) and Ovx+E rats (significant dose x time interaction, $F_{168.0, 1195} = 1.384$, p=0.0017) following administration of OLZ (**A,B**). Ovx rats experienced significant reductions in locomotor activity relative to vehicle at 10 and 30 min (0.3 mg/kg) and 230 min (0.3 and 1.8 mg/kg) post-dosing. Ovx +E rats experienced significant reductions relative to vehicle at 170 min (0.56 mg/kg) post-dosing. p<0.05, horizontal colored lines matching the respective dose color represent the 10-minute bins at which OLZ treatment was significantly different from vehicle treatment. n=7-9 rats per group.
